# Impact of Holocene environmental change on the evolutionary ecology of an Arctic top predator

**DOI:** 10.1101/2022.10.06.511126

**Authors:** Michael V. Westbury, Stuart C. Brown, Julie Lorenzen, Stuart O’Neill, Michael B. Scott, Julia McCuaig, Christina Cheung, Edward Armstrong, Paul J. Valdes, Jose Alfredo Samaniego Castruita, Andrea A. Cabrera, Stine Keibel Blom, Rune Dietz, Christian Sonne, Marie Louis, Anders Galatius, Damien A. Fordham, Sofia Ribeiro, Paul Szpak, Eline D. Lorenzen

## Abstract

The Arctic is among the most climatically sensitive environments on Earth, and the disappearance of multiyear sea-ice in the Arctic Ocean is predicted within decades. As apex predators, polar bears are sentinel species for addressing the impact of environmental variability on Arctic marine ecosystems. By integrating genomics, isotopic analysis, morphometrics, and ecological modelling, we investigate how Holocene environmental changes affected the evolutionary ecology of polar bears around Greenland. We show that throughout the last ∼11,000 years, Greenlandic polar bears have been heavily influenced by changes in sea-surface temperature (SST) and sea-ice cover. Most notable are major reductions in effective population size at the beginning of the Holocene and during the Holocene Thermal Maximum ∼6 kya, which coincide with increases in annual mean SST, reduction in sea-ice covers, declines in suitable habitat, and shifts in suitable habitat northwards. Furthermore, we show how individuals sampled from west and east Greenland are genetically, morphologically, and ecologically distinct. We find bears sampled in west Greenland to be larger, more genetically diverse and have diets dominated by ringed seals, whereas bears from east Greenland are smaller and less diverse with more varied diets, putatively driven by regional biotic differences. Taken together, we provide novel insights into the vulnerability of polar bears to environmental change, and how the Arctic marine ecosystem plays a vital role in shaping the evolutionary and ecological trajectories of its inhabitants.

**Teaser:** Multivariate investigations of the environment’s role in the evolutionary ecology of Greenlandic polar bears.

## Introduction

The Arctic is among the most climatically sensitive environments on Earth, with increases in temperatures being two- to three-fold higher at the northern latitudes than the global mean (*1, 2*). Within decades, increasing temperatures resulting from Arctic amplification of global warming are expected to lead to the disappearance of multiyear sea-ice in the Arctic Ocean (*2*). Such changes in temperature and sea-ice cover represent thresholds in the Earth’s climate system, potentially leading to irreversible ecosystem change (*3, 4*).

Although the Arctic is currently experiencing drastic alterations associated with anthropogenic climate change (*5*), past environmental and ecological changes linked to climatic shifts in the region are well preserved in palaeo-archives (*6–8*). Palaeo-archives reveal, for example, the impact of the Greenland Ice Sheet on the biogeochemistry and productivity of surrounding marine ecosystems (*9, 10*). These Greenlandic marine ecosystems are further influenced by sea-ice cover, sea-level fluctuations, changes in ocean circulation, and the availability of nutrients and light for biological productivity, among others (*11*). The impact of such changes on the evolutionary ecology of Arctic species are yet to be fully explored.

Changes in environmental conditions can greatly impact the base of the Arctic food web (*12*). These shifts at the base of the food web drive bottom-up changes in ecosystem structure and function by altering pelagic secondary production, the main food source for all higher trophic-level organisms (*13*). Shifts in food sources can influence the mobility and connectivity of species (*14*) and disrupt gene flow among populations, altering evolutionary pathways (*15*). Nearly all organisms within an ecological community are directly or indirectly tied to the top of the food chain (*16*). Therefore, an approach to assess the impacts of environmental change on ecosystems is to investigate apex predators, providing top-down insights into the ecosystem as a whole (*17*). In the Arctic marine realm, polar bears (*Ursus maritimus*) occupy the top predatory niche. They are found on ice-covered waters across the region, and depend on sea-ice primarily for hunting pinnipeds (*18*). While polar bears have been found to inhabit multiyear pack ice in the central Arctic basin, their preferred habitat is seasonal sea-ice, including land-fast ice, around the coastline of the Arctic Ocean region (*19*). This reflects the exploitation of relatively higher levels of biological productivity around seasonal sea-ice by seal species (*20*).

Polar bears have evolved a novel and distinct ecology, behaviour, and morphology in response to life in the high Arctic (*21*). The species is split into 19 management units, determined from a combination of traditional knowledge, radiotelemetry, movements of adult females with satellite radio-collars, and genetics (*22–24*). Greenland is home to at least three management units: Kane Basin and Baffin Bay on the west coast, and East Greenland on the east coast (*25*). A distinct and previously unknown subpopulation was recently documented in the southern coast east Greenland (*26*), highlighting that much is still unknown about the complexity of polar bear populations around Greenland.

The dependence of polar bears on sea-ice makes them a sentinel species for detecting ecosystem tipping points (*18*). To assess the vulnerability of polar bears to past and future climate and environmental change, it is vital to apply a complementary suite of approaches that combines insights from the present with those of the past (*7, 27*). Population genomics provide insights into population subdivision and the longer-term demographic history of polar bear populations (*21, 28*). However, it is difficult to elucidate the drivers underpinning these changes using genetics alone. Dietary tracer analysis of stable carbon (*δ*^13^C) and nitrogen (*δ*^15^N) isotope data and ecological models can provide important additional information for determining biotic and abiotic drivers of past demographic change (*29, 30*). Stable isotopes can elucidate current population-specific foraging behaviour and regional primary productivity (*31*), while ecological models can be used to reveal shifts in the size and area of ecologically suitable habitats in response to climate and environmental changes (*7, 30*). The incorporation of morphological data can be used to consolidate evolutionary and ecological insights, as changes in morphological phenotype can be shaped by both genetics and the environment (*32*).

Here, we aim to better characterise the current genomic, morphometric, and dietary relationships among polar bears around Greenland, as well as the role the environment played in shaping their evolutionary ecologies and distributions throughout the Holocene. We focus on integrating independent yet complementary datasets and analyses for polar bears sampled from around Greenland (Fig 1A), and make inferences about their past evolutionary history and the ecological drivers of similarities and differences between modern bears inhabiting the west and east coasts.

**Figure 1:**
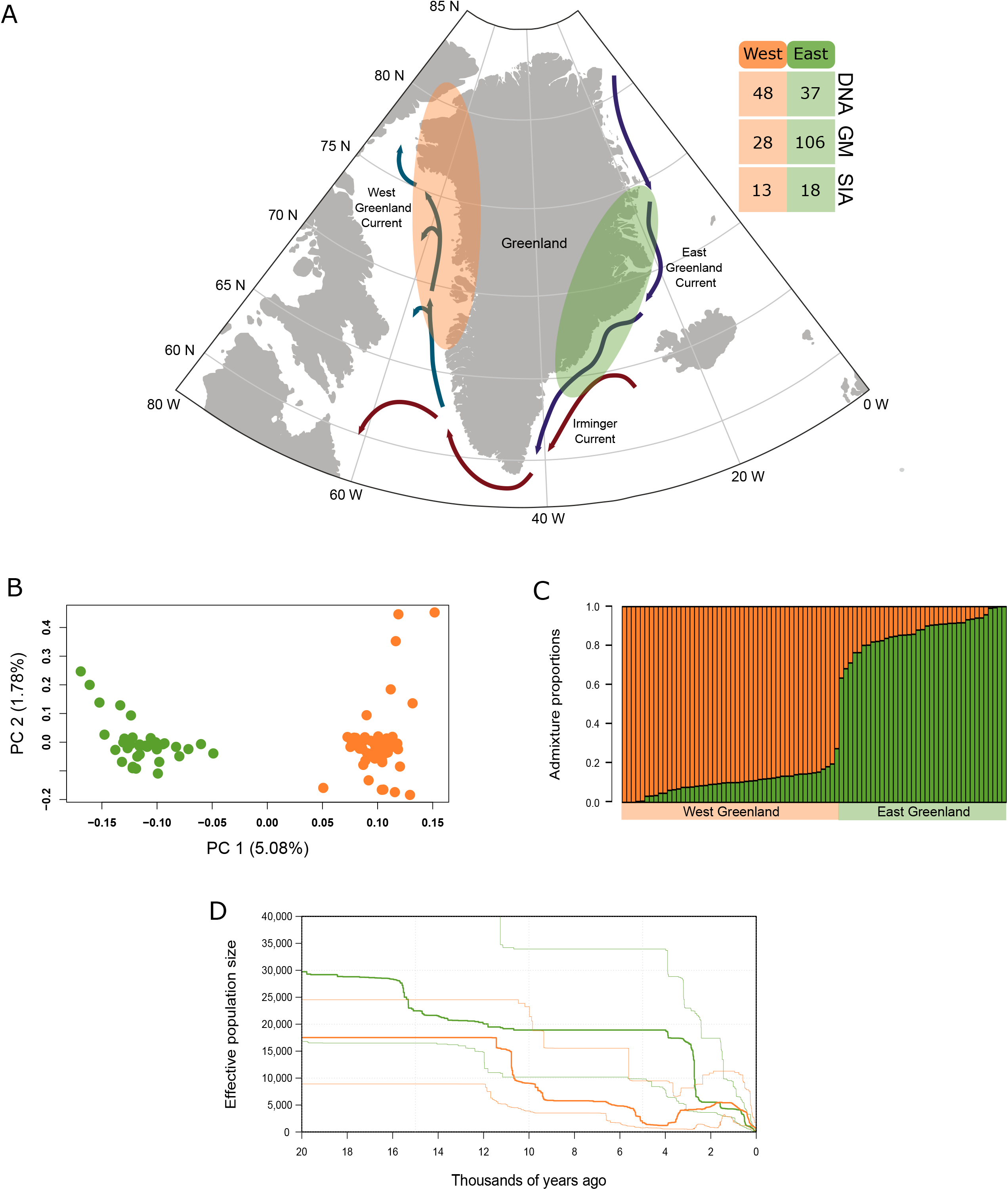
(A) Map of the study area. Shaded areas show the approximate spatial area of sampling locations of the polar bear specimens analysed from the west (orange) and east (green) coasts of Greenland. Polar bear sample sizes are indicated for population genomics (DNA), morphometrics (GM), and stable isotopes (SIA). The study also included genomic data from polar bears sampled in adjoining areas in Canada and Svalbard, and *δ*^13^C and *δ*^15^N stable isotope data from the five predominant pinniped prey species of polar bears (bearded seal, harp seal, hooded seal, ringed seal, and walrus), see Materials and Methods for further details. Arrows show ocean currents, modified after (*128, 129*). Genomic population structure analyses based on 85 Greenland polar bears; (B) principal component analysis, with the percentage of variance explained by each component indicated between parentheses, and (C) admixture proportion analysis. (D) Changes in effective population size (*N*_*e*_) through time inferred using individuals with genome-wide coverages >20x (west n = 12, east n = 9) using the site frequency spectrum and stairway plots. Thick lines show the median *N*_*e*_ values and the faded lines show the 97.5% confidence intervals.

## Results

### Genomics

After mapping to the polar bear reference genome, we obtained 106 polar bear nuclear genomes ranging in coverage from 1.7x - 32.8x (including a single 114.1x individual) (Supplementary Table S1). Population structure analyses (PCA and admixture proportions) showed clear differentiation between polar bears sampled from the west (n=48) and east (n=37) coasts of Greenland (Fig. 1B, C, and Supplementary Fig. S1). We did not observe genetic structuring between polar bears sampled from the Kane Basin and Baffin Bay management units of the west coast (Supplementary Fig. S1), in accordance with previous results (*22*). We found a closer affinity of bears from the west coast to Canadian individuals, and of bears from the east coast to the neighbouring individuals from Svalbard (Supplementary Fig. S1). The fixation index (*F*_ST_) between polar bears sampled from the west and east Greenland coasts was significant (0.0174 ; p-value < 2.2e-16), further supporting clear genomic differentiation between the two coastlines.

We found similar levels of heterozygosity across all individuals, with overlapping 95% confidence intervals. The higher mean heterozygosity observed in west bears (0.000831) compared to east bears (0.000773) (Supplementary Table S2) was significant (*t* = 2.6804, *p* = 0.01985). We did not observe significantly higher mean levels of nucleotide diversity in west Greenlandic bears (0.00033) compared to the east (0.00033) (*t* = 0.391, *p* =0.6961). We found three individuals with inbreeding coefficients (probability of two alleles being identical by descent, F) greater than 1%; all three individuals were from the east coast (PB_9 F=26.8%, PB_28 F=3.8%, and D24082 F=2.8%)

Estimates of the deeper (>20 thousand years ago [kya]) demographic history of the bears using the pairwise sequential Markovian coalescent (PSMC) model (*33*) yielded identical demographic trajectories for all individuals analysed, regardless of their geographic origin or management unit (Supplementary Fig S2). Results showed relatively stable effective population sizes (*N*_*e*_) from 150 kya until ∼50 kya, after which the joint population experienced a decline in *N*_*e*_ culminating in an approximately 50% decline in *N*_*e*_ by ∼20 kya. The PSMC results indicated an increase in *N*_*e*_ ∼15 kya, but the reliability of the PSMC model to infer changes in *N*_*e*_ within the last 20 thousand years is negligible (*33*).

Estimates of the more recent (<20 kya) demographic dynamics using stairway plots (*34*) revealed similar demographic trajectories in the west and east groups, with overlapping confidence intervals (Fig. 1D). In bears from both coasts we see overall declines in *N*_*e*_ over the last ∼15,000 years, caused by several rapid declines throughout. We investigated the impact of sample sizes and found the results may lose some of the more fine-scale population size changes when based on fewer individuals, but with overall similar trajectories (Supplementary Fig. S3). Hence, we may be missing some more fine-scale inferences in the east Greenland bears (n = 9) relative to west Greenland (n =12). Results were similar when specifying the giant panda, instead of the spectacled bear, as the ancestral state but with more rigid changes in *N*_*e*_ and therefore less resolution overall (Supplementary Fig. S4).

Calculations of genome-wide Tajima’s D values uncovered positive values for both west (mean value of 0.196) and east (mean value of 0.030) Greenland bears. A positive score signifies low levels of both low- and high-frequency polymorphisms, and may reflect recent population contractions (*35*). However, only west Greenland bears showed a significant deviation from 0 (west p=0.0003, east p=0.2837).

### Stable isotopes

Our *δ*^13^C (feeding habitat) and *δ*^15^N (trophic level) data from 31 polar bears comprising seven bears sampled from the Kane Basin management unit, six bears from the Baffin bay management unit (both west Greenland), and 18 bears sampled from the east Greenland management unit, and their predominant pinniped prey species sampled from the two coasts (n = 110; west =54, east= 56) (Supplementary Table S3), enabled us to assess how the taxonomic composition of diets compared between coasts. After accounting for a decline in *δ*^13^C values caused by the burning of fossil fuels (the Suess effect), bears from the west coast of Greenland had significantly higher *δ*^13^C (*U* = 14, *p* < 0.001) and *δ*^15^N values (*t* = 8.10, *p* < 0.001), relative to bears sampled from the east coast of Greenland (Fig. 2A and 2B). Although west coast bears were sampled from two adjoining management units, we did not observe any significant differentiation in their isotopic compositions (p>0.05) (Supplementary Fig. S5).

**Figure 2.**
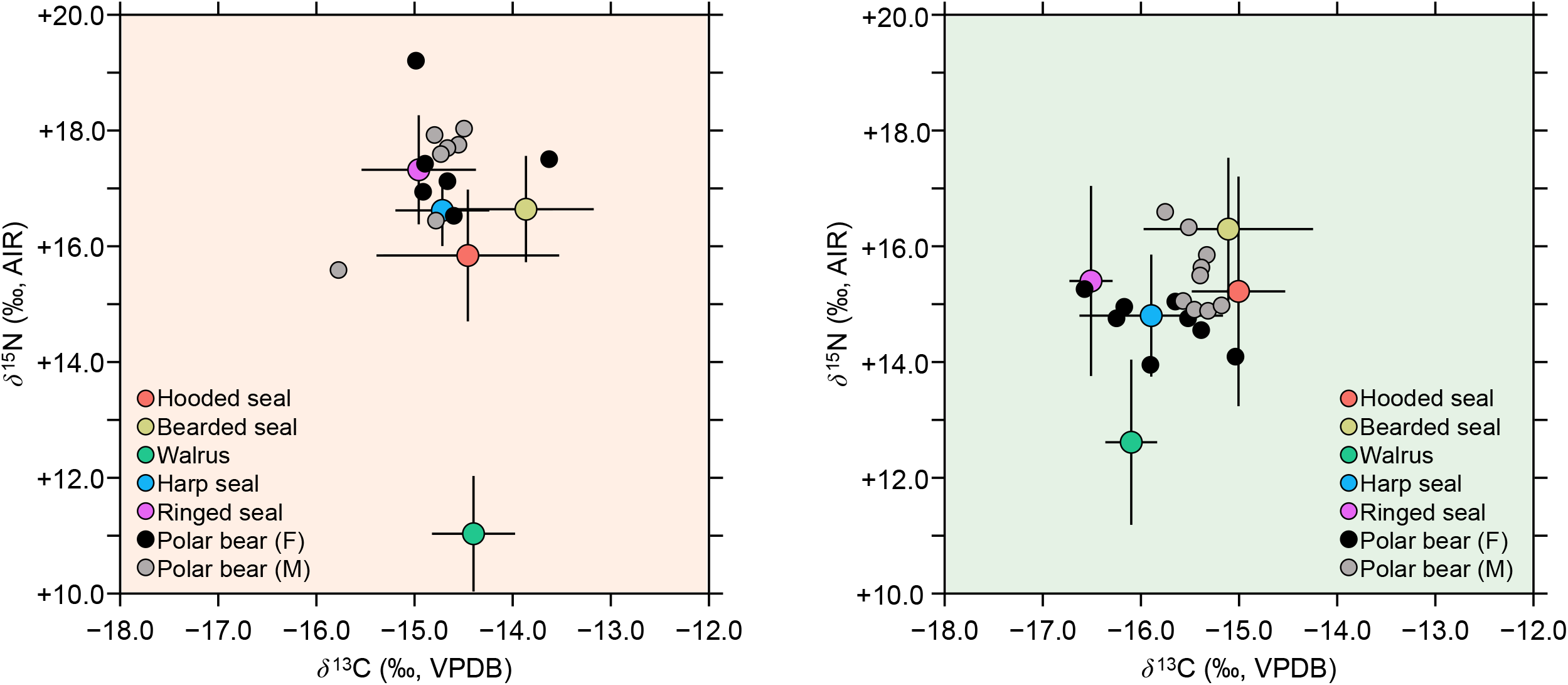
Comparison of Greenland polar bears and five potential prey bone collagen *δ*^13^C and *δ*^15^N values; (A) west coast, (B) east coast. Polar bears have been adjusted by −0.5 ‰for δ^13^C and −4.0 ‰for *δ*^15^N to account for the offset between predator and prey.

A qualitative comparison of the polar bear and prey isotopic compositions, accounting for trophic discrimination, revealed that bears sampled in west Greenland clustered most closely with ringed seals, suggesting a diet dominated by this species (Fig. 2A), in agreement with previous work (*36*). In contrast, bears from the east coast had more varied diets. However, they qualitatively clustered most closely with harp seals and bearded seals, indicating their primary prey was harp seal, followed by bearded seal (Fig. 2B).

Within bears from the west, there were no significant sex-based differences in *δ*^13^C (*U* = 20, *p* = 0.94) or *δ*^15^N (*U* = 17, *p* = 0.62). Although males from the east had higher *δ*^13^C values than females, this difference was not statistically significant (*t* = 2.06, *p* = 0.07). Males from the east had significantly higher *δ*^15^N (*U* = 11.5, *p* = 0.01) values relative to females from the same area. The higher *δ*^13^C and significantly higher *δ*^15^N values in the male relative to female polar bears in the east (Fig. 2B) suggest that females consumed primarily ringed seals and harp seals, while males consumed much larger amounts of bearded seals and hooded seals. A similar pattern was not observed in polar bears in the west.

### Geometric morphometrics (GM)

Investigations into shape differences revealed highly significant shape differences (*p*<0.0001) between polar bear individuals sampled in the west (from both Kane Basin and Baffin Bay management units) and east (from the East Greenland management unit). Despite being from different management units, bears sampled on the west coast did not show significant differences, and so were pooled together. Relative to skulls sampled in the east, skulls from bears sampled in west Greenland were slightly narrower posteriorly, and had a temporal fossa that was extended dorsoposteriorly, but anteriorly truncated, an orbit that was slightly compressed dorsally, while the anterior point of the premaxilla was slightly elevated (Fig. 3A, B). Moreover, 82.1% of the skulls could be correctly reclassified to sampling coast (west *vs* east) by shape after cross validation (Table 1). Skulls tended to be largest (centroid) for both sexes in the individuals from the west. However, this difference was only statistically significant for females (p=0.006), but not for males (p=0.077) (Fig. 3C).

**Table 1:**
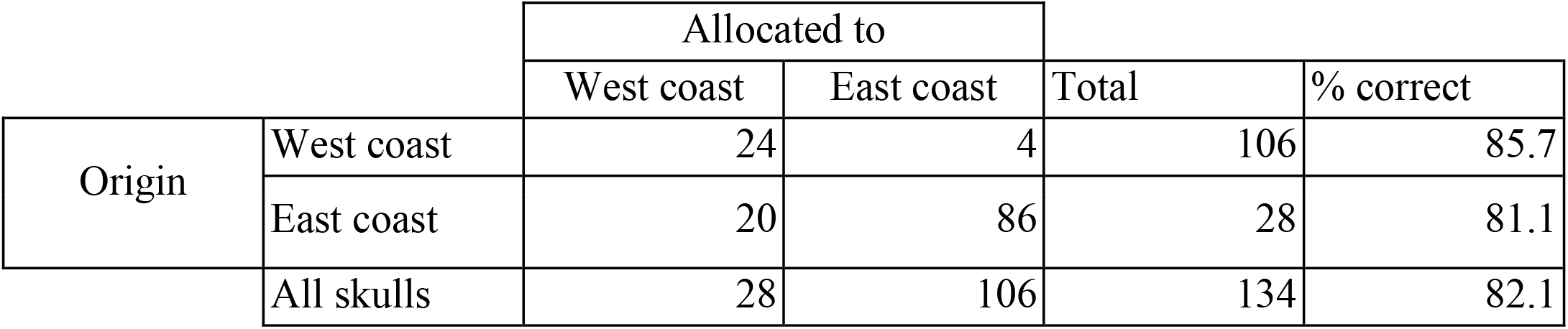
Success rate of reclassification of specimens to geographical area by skull shape using jackknife cross-validation.

**Figure 3:**
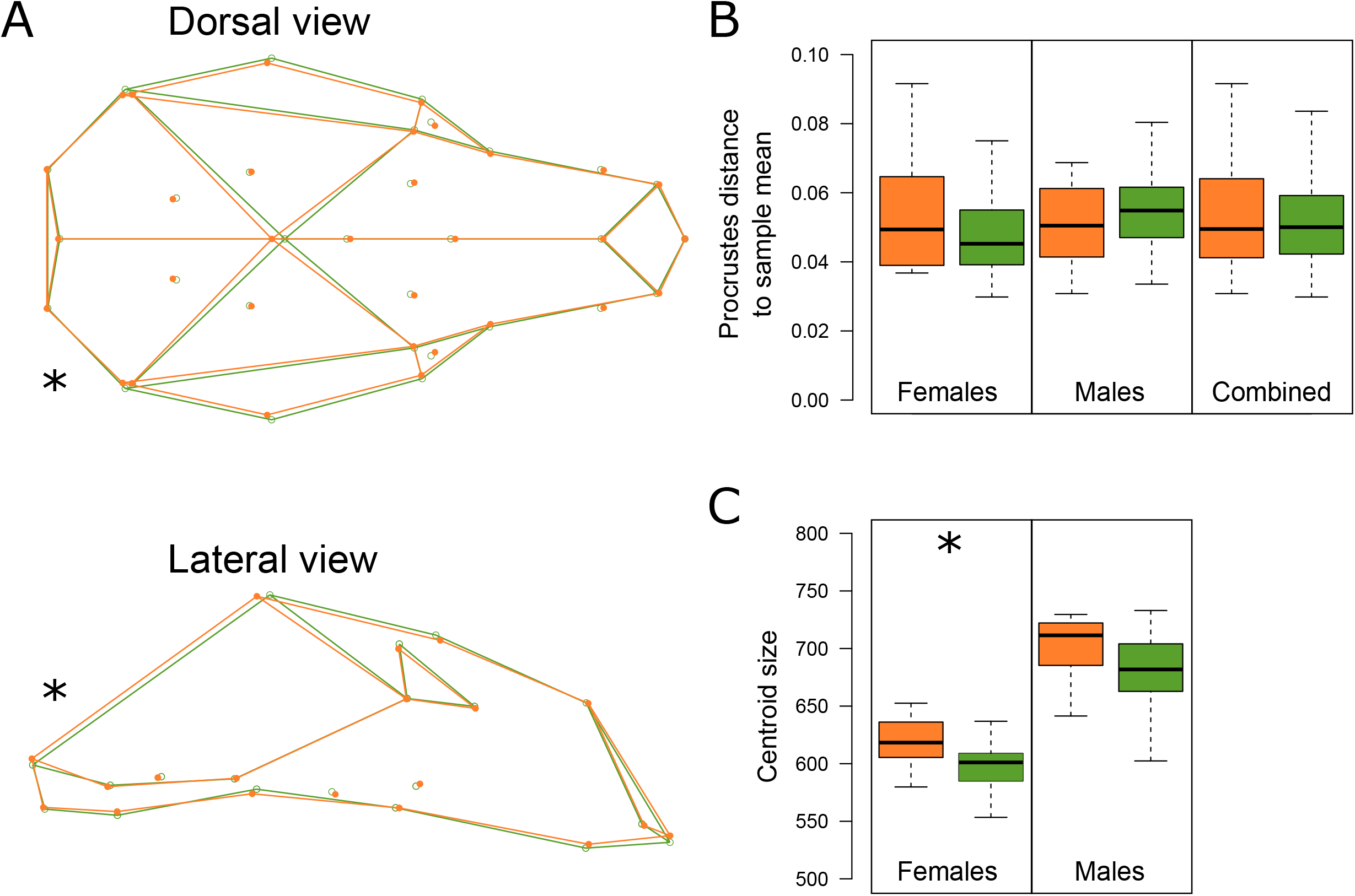
Geometric morphometric comparisons of 134 polar bears sampled in west (orange, n = 28) and east (green, n = 106) Greenland. (A) Mean area-specific shapes of polar bear skulls in dorsal and lateral aspects. Correction for allometric effects was performed (see Materials and Methods for further details). (B) Procrustes distances of specimens to the mean of the relevant sample of the whole dataset combined and separated by sex. (C) Centroid sizes (square root of sum of squared distances of landmark positions to configuration centroid) of skulls from male and female polar bears from the west and east coasts of Greenland. Asterisks indicate significant differences between west and east Greenland bears.

### Habitat suitability

Ecological niche modelling (ENM) tuning and cross-validation resulted in the optimised model having high geographic transferability shown by low differences between spatial cross validation folds according to the Area Under the Receiver Operating Curve values (AUC_diff_ = 0.05), low 10^th^ percentile training omission rates (OR10 = 0.13), and moderate AUC values (0.75 ± 0.05). The model had a greater regularisation multiplier (i.e. complexity penalisation; RM = 3) and fewer feature classes (FC = 3) than the MaxEnt default (RM = 1 and FC = 4). The ENM had good ability to discriminate between areas of high suitability at known occurrences and background sites for the contemporary period that the model was built on (Boyce index = 0.82; Supplementary Fig. S6)

Our ENM revealed increasing probability of occurrence (hereafter habitat suitability (*37*)) of polar bears at sea-surface temperatures (SST) between 0°C and ≲ 4.5°C (Supplementary Fig. S7), declining to 0 suitability by SST of 15°C. Habitat suitability was positively correlated with annual average sea-ice concentration, and decreased slightly with increasing sea-surface salinity. Habitat suitability was not affected by changes in annual variation in fractional sea-ice cover or annual variation in SST (Supplementary Fig. S7). Habitat suitability was high at the beginning of the Holocene (∼11 kya) when areal average SST was around 1.5 °C, declining sharply with warmer SST values and lower annual average sea-ice concentrations (Fig. 4A, B and Supplementary Figs. S8 and S9). The ENM projected a decreasing trend in average habitat suitability from 11 kya to 4.5 kya, whereafter mean habitat suitability values fluctuated around a long-term stable average (Fig. 4C). Analysis of the centre of gravity of habitat suitability, indicated a northward shift in the highest habitat suitability values which corresponded to declines in areal habitat suitability (Fig. 4D). All predictor variables show similar patterns to habitat suitability with the exception of SST seasonality, which decreased steadily from a peak ∼7.5 kya (Supplementary Fig. S9). Animations of habitat suitability through time for the study region are provided as a supplementary video (Supplementary Video S1).

**Figure 4:**
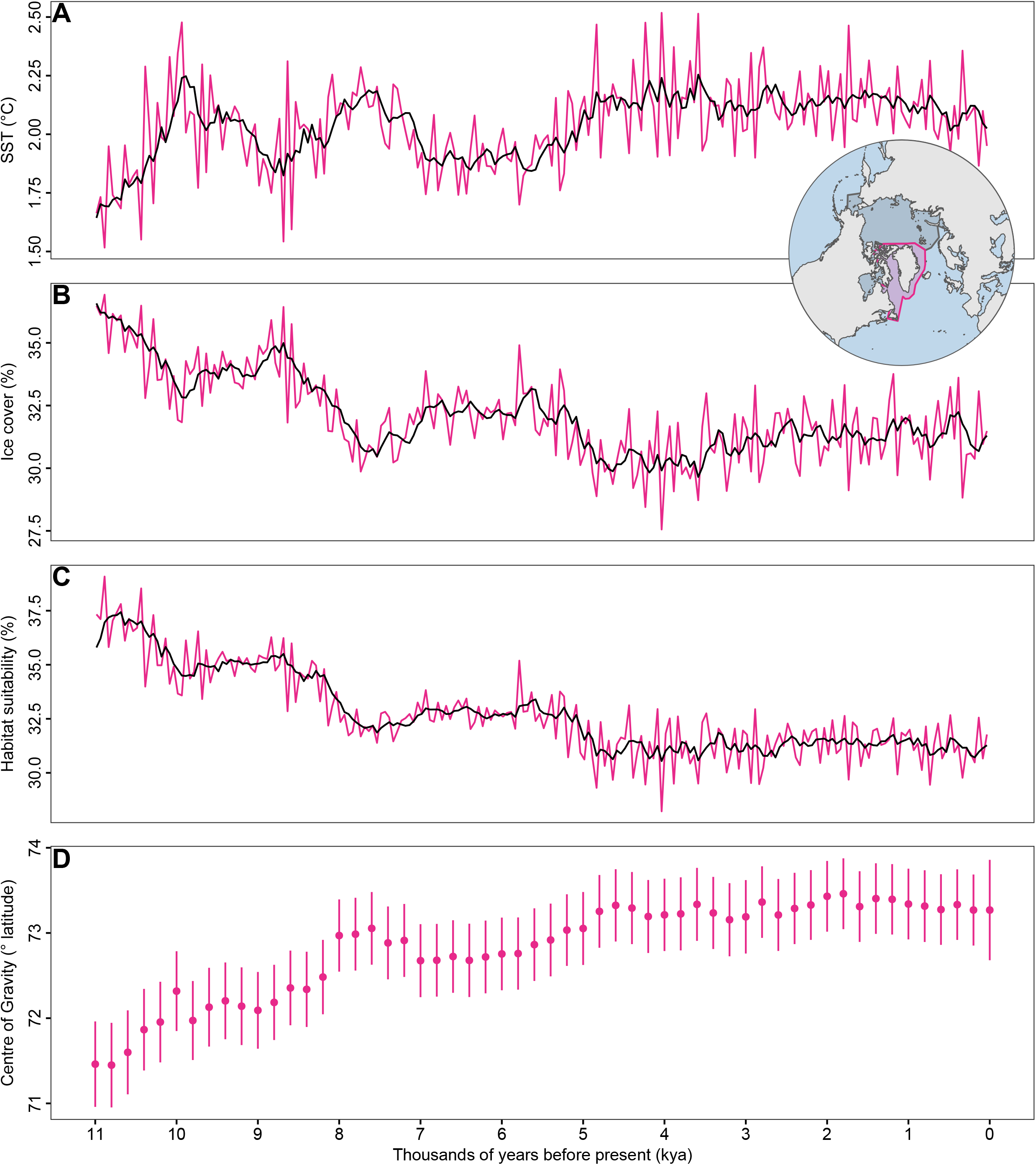
Areal means of the variables used in the model and predicted habitat suitability from 11 kya to 0 kya. The panels show the areal mean values in sea surface temperature (SST), sea-ice cover, and mean habitat suitability for the Greenland study region. Red lines in each plot show 1,000 year smoothed mean values. The inset map shows the circumpolar region in dark blue and the Greenland sub region in red.

## Discussion

Using an integrative multi-proxy approach, we reveal similar evolutionary trajectories, but distinct ecologies, of polar bears sampled from the west and east coasts of Greenland. In doing so, we can form hypotheses about the abiotic and biotic factors that have differentially impacted the connectivity, demographic history, diet, and morphology of polar bears around Greenland.

An association between *N*_*e*_ and changes in environment is visible when comparing our genomic demographic reconstruction of the past 20 kya (Fig. 1D) with palaeoclimate and suitable habitat over the same time period (Fig 4). Overall, we observe a clear pattern of *N*_*e*_ decline when suitable habitat decreased, which was associated with periods of warmer sea-surface temperature (SST) and reduced sea-ice cover. SST and sea-ice cover are highly correlated, with the latter more likely to be the causative factor driving declines in polar bears, which rely on sea-ice for movement, mating, and hunting (*38*). In fact, there is evidence from contemporary populations of declines in polar bear survival and body condition associated with sea-ice loss (*39–41*), and increased overall vulnerability of modern populations has been predicted as sea-ice continues to decrease in coming generations (*42*).

Although associated with large confidence intervals, the major and rapid decline in *N*_*e*_ in west Greenland bears at the onset of the Holocene ∼11 kya (Fig. 1D) coincided with a decline in range-wide mean suitable habitat of ∼5%, an increase in range-wide mean sea surface temperature (SST) of ∼0.25 °C, and a reduction in range-wide mean annual sea-ice cover of ∼6% (Fig. 4). At the same time, we observed a coincident shift in the centre of gravity of habitat suitability to higher latitudes, suggesting the decrease in suitable habitat caused a northwards shift of the bears, potentially driven by the decreasing annual sea-ice cover. We do not observe a coinciding decline in median *N*_*e*_ in the east Greenland bears, although the 97.5% confidence intervals do indicate decreasing *N*_*e*_ at ∼11 kya (Fig. 1D). The lack of concordance between the east Greenland bears and climate could reflect a smaller dataset (12 bears in west *vs* 9 bears in east); we show estimates of population fluctuations may become artificially older when fewer individuals were included in the analysis (Supplementary Fig S3).

Another major and rapid decline in *N*_*e*_ in polar bears from the west coast beginning ∼6 kya coincided broadly with the Holocene Thermal Maximum (8 kya - 5 kya (*43*)), a decline in habitat suitability of ∼4%, an increase in range-wide mean SST of ∼0.5 °C, and a reduction in annual range-wide average sea-ice cover of ∼6% (Fig 4). These align with a shift in the centre of gravity of habitat suitability to higher latitudes, again suggesting the decreases in suitable habitat may be caused by a loss of suitable habitat in the south of the range. In contrast, we do not see a decline in *N*_*e*_ in polar bears from east Greenland at the same time period, which may again reflect smaller sample size for the analysis and therefore lower resolution.

A decline in *N*_*e*_ may reflect either decreasing census population size, or loss of population connectivity. The latter was not supported by our habitat models. We did not observe any increase in fragmentation of suitable habitat at these times (Supplementary Video S1) and therefore propose the observed major declines at both 11 kya and again at 6 kya were driven by a decrease in census population size.

The association between declines in *N*_*e*_ and corresponding increases in SST and decreases in sea-ice cover across the Holocene suggests that ongoing climate change could have a major detrimental effect on polar bear populations around Greenland. Simulated ocean warming patterns under a range of different future climate scenarios show extensive SST warming around Greenland (90% CI SSP1-2.6 = -0.8 | 2.4 °C; 90% CI SSP5-8.5 =0.6 | 5.4 °C, above 1981-2010 baseline) and reductions in annual sea-ice extent by the end of the century (90% CI SSP1-2.6 = - 17.3 | -1.9%; 90% CI SSP5-8.5 = -32.8 | -13.7 %, from 1981-2010 baseline), even under reduced emission scenarios (*44*). It is expected that the Arctic will become essentially ice-free in summer before the end of the 21st century, but is very likely to remain sea-ice covered during winter for all warming scenarios (*45*).

Although we observed more recent fluctuations in *N*_*e*_ fluctuations over the past 3,000 years, with sharp declines in east Greenland ∼3 kya and on both coasts from ∼1 kya, we did not observe any notable concurrent changes in suitable habitat, nor in the individual palaeoclimate proxies used in our analysis (including SST, sea-ice cover, salinity) (Fig 4A, B, Supplementary Fig. S8). This may reflect the temporal resolution of our climate data (*46*), which is based on long-term climatic means that have been shown to affect ecological responses to environmental and climatic variables (*47*). It is noticeable that the recent population declines beginning ∼1,000 years ago coincide with the Medieval climate anomaly, a period in which marine and lake multi-proxy records suggest the North Atlantic was atypically warm (*6*). Similar to our findings at the end of the Holocone and during the Holocene Thermal Maximum, this further supports that increased temperatures and concurrent declines in sea-ice may have a detrimental effect on polar bear population size and/or connectivity.

The strikingly similar patterns of demographic change throughout the past 150,000 years observed in our PSMC results, suggest bears from the two coasts either responded in identical manners to past environmental changes, or they comprised a single evolutionary unit and were likely panmictic until the very recent past (Supplementary Fig S2). The latter is congruent with our analyses of population subdivision that showed significant, albeit low (*F*_ST_ = 0.0174), levels of genomic differentiation between bears sampled from west and east Greenland, with some degree of admixture (Figs. 1C). Differentiation between coasts was further documented by our dietary and morphological analyses of the bears, which indicated distinct foraging ecologies with significant differences in skull shape and size based on locality (Figs. 2 and 3). Our findings across datasets (genetic, stable isotopes, and morphometrics) indicate clear differentiation and a putative ecologically-driven genetic divide, rather than through longer-term evolutionary forces.

Further supporting a recent divergence is our finding of consistent and continuous suitable habitat over the north of Greenland throughout the Holocene (Supplementary video S1), which is likely to have facilitated movement and connectivity between west and east. The ability of polar bears to migrate vast distances over the northern Arctic has been demonstrated using telemetry tracking; a single female moved from Kane Basin (west coast) northwards around Greenland to Russia - a distance of ∼2,000 kilometres (*48*). Our habitat models are based exclusively on coarse-scale abiotic factors, and thus assume no demographic or local-scale geographic barriers. Therefore, the observed differentiation between west and east coast bears may well reflect biotic factors. Our stable isotope ecological proxy data support this, and reveal ecological distinctions between bears sampled from each coast (Fig 2). Different ocean currents result in a longer open-water season and higher productivity along the west Greenland coast, and a much less productive marine environment along the east coast, where sea-ice cover is more extensive in both time and space (*49*) (Fig. 1A). These differing ocean currents and levels of productivity could be the ecological driver, as variation at the base of food webs have bottom-up effects on ecosystem structure and function as a whole (*13*). In addition, our stable isotope analysis of five pinniped species (Fig 2) revealed that differences between geographic localities reflect both discrepancies in the type of prey species consumed, and in the underlying primary producers, i.e. relative abundance of sea-ice microalgae and phytoplankton, as has been shown in other Arctic predators (*31*).

The dietary plasticity of polar bears indicated by our findings may be a key attribute in the resilience of the species, acting to buffer the relatively low levels of genetic variation for selection to act upon (*21, 28, 50*). Prey abundances are much lower in east Greenland and therefore bears in this region may be especially prone to dietary shifts (*51*). Indeed, dietary shifts in east Greenland polar bears due to sea-ice reduction in recent decades have been reported, with a decline in ringed seal consumption and an increase in harp seals and hooded seals (*52, 53*); a finding supported by our data. In addition, polar bear population density may also play an important role. However, we are currently limited in our ability to elucidate this further, as census size estimates for the East Greenland management unit are unavailable (*51*).

The tendency for larger skull size, and by proxy larger body size/mass (*54*), in bears sampled in west Greenland (Fig. 3) may reflect underlying genetic mechanisms. Alternatively, differences may be driven by phenotypic plasticity and adaptation to the unique diets on each coast. The ability of west Greenland bears to grow larger than their east Greenland counterparts may reflect the relatively higher amounts and/or more nutrient-rich food in the more productive west (*49*). In east Greenland, size differences between sexes may have driven additional ecological divergence; higher *δ*^13^C and *δ*^15^N values in males (Fig. 2B) indicated the incorporation of larger amounts of bearded seals in their diets, while females consumed primarily ringed seals and harp seals. Larger males have a greater capacity to successfully hunt larger prey than smaller females, and bearded and hooded seals are larger (*55*). Therefore, due to a decline in the availability of smaller prey, east Greenland males may hunt larger prey. This is supported by their wider posterior skulls (Fig. 3A) - particularly around the temporal fossa, where the *temporalis* muscle that operates the mandible attaches, suggesting more muscular heads and adaptation to larger prey species. If ringed seals are the optimal polar bear prey species in terms of energetic return for energy input, the finding in west Greenland may be driven by higher productivity and greater prey availability, allowing both males and females to converge on the same diet (*56*).

Taken together, our results suggest that the size and connectivity of polar bear populations around Greenland have been influenced by past changes in sea surface temperature and sea-ice cover. Our findings also suggest bears may ecologically adapt relatively rapidly. This ability is crucial for resilience to ongoing rapid shifts in the distributions and abundances of prey caused by anthropogenic climate change (*57*). Adaptation through phenotypic and/or ecological shifts is more rapid than by DNA nucleotide changes, and is a major factor allowing species to survive conditions of changing climate and food availability (*58*). However, as we find that declines in overall suitable habitat are driven by a loss in the southern parts of the habitat, even with behavioural and phenotypic plasticity, the pace and scale of predicted near-future ecological and environmental change in the Arctic (*45*), coupled with a long generation time, is likely to leave polar bears vulnerable to climate and environmental change in the near future.

## Materials and Methods

### Samples

For the genomic analysis, we utilised publicly available raw Illumina fastq files from available polar bear individuals sampled across the Atlantic Arctic (*21, 26, 28, 59*). The data included 48 samples collected from west Greenland, including the management units of Kane Basin (n = 27) and Baffin Bay (n = 18) and 37 samples collected from east Greenland, which is currently recognised as a single management unit (*21*). Furthermore, we included available genomic data from polar bears of the adjoining subpopulations on either side of Greenland: Canada (*59*) and Svalbard (*28*). All data is available from NCBI under the following Bioproject IDs: PRJNA169236, PRJNA196978, PRJNA210951, and PRJNA669153.

For *δ*^13^C and *δ*^15^N stable isotope analyses we sampled bone powder from a total of 31 polar bears from west Greenland (Kane Basin n = 7; Baffin Bay n = 6) and east Greenland (n = 18) (Supplementary Table S3). To control for region-specific differences in primary productivity, we also performed stable isotope measurements on the bone collagen of the five main putative pinniped prey species of polar bears (*20, 60*): bearded seals, harp seals, hooded seals, ringed seals, and walruses. The total of 110 pinniped specimens analysed represented 54 individuals from the west and 56 individuals from the east (Supplementary Table S3). The prey species were sampled across multiple decades and therefore there may be subtle variations in the isotopic compositions of each of these, as has been observed in other taxa (*61*). Nevertheless, we think that these data are suitable for our purposes for two reasons. First, while there may be some subtle variations in the isotopic compositions of the prey over multiple decades, this variation will tend to be minimised since we sample bone collagen from these prey species, a tissue with a very slow rate of turnover that will tend to dampen this variation and average the diet over the life of the individual (*62*). Second, we have avoided attempting a more precise, quantitative assessment of polar bear diet composition using a stable isotope mixing model because of these uncertainties about the isotopic compositions of these prey. This more qualitative approach is more conservative in that we limit our interpretation to relative, rather than quantitative, reconstructions of polar bear diets.

For geometric morphometrics we analysed a total of 134 specimens; 28 were sampled from the west coast of Greenland (Kane Basin n = 14; Baffin Bay n = 14, 14 males and 14 females) and 106 from the east coast (58 males and 48 females) (Supplementary Table S4). The skulls were sampled between 1936 and 2015 and we make the assumption that skull size and shape did not change through this period.

There was partial overlap between the samples used for the different analyses. There was one sample that overlapped in both the genomic and stable isotope analyses, six samples overlapped between the genomic and geometric morphometric analyses, and 26 samples that overlapped between stable isotope and geometric morphometric analyses.

For stable isotope analysis and geometric morphometrics, only specimens representing adult individuals were sampled. For stable isotopes, this was to avoid biases that could occur in juveniles and sub-adults, as higher nitrogen isotope values are found in young individuals feeding on their mother’s milk (*63, 64*). For geometric morphometrics, this was to avoid excessive allometric variation in the sample. Adult males were defined when having at least six complete dentin or cementum growth-layer groups (equal to ≥ 6 years of age), and adult females when having at least five growth-layer groups (equal to ≥ 5 years), using the methods of (*65*). In brief, the age determination followed the procedure of counting annual growth-layer groups in the cementum of the lower right I3 using decalcification, thin sectioning (14 µm), and staining (*66*).

## Genomic analyses

### Raw data processing

We processed all raw sequencing reads with the PALEOMIX (*67*) pipeline. Internally, adapters, stretches of Ns, and low-quality bases in reads were trimmed and filtered with AdapterRemovalv2 (*68*) using default parameters. BWA-backtrack v0.7.15 (*69*) was used to map the cleaned reads to the polar bear pseudo-chromosome genome (*21, 50*) (based on Genbank accession: GCA_000687225.1), mitochondrial genome (Genbank accession: NC003428) included, with default parameters. Reads with mapping quality < 30 were filtered using SAMtools (*70*). Duplicates were removed with Picard v2.6.0 (*71*). Possible paralogs were filtered using SAMtools v1.6. Finally, local realignment around indels was done using GATK (v 3.3) (*72*).

### Population genomics

To determine the level of genome-wide population structure among Atlantic Arctic polar bears, we performed multiple principal component analyses (PCA). As our dataset contained both high- and low-coverage genomes, we first sought to investigate whether different sequencing coverage of individuals influences results. To do this, we ran a PCA using all 85 Greenlandic polar bears. To put the population structure into a regional perspective, we also ran a PCA with (iii) all 85 Greenlandic samples, and individuals sampled from the surrounding regions to the west (Canada, n=3) and further east (Svalbard n= 18)).

As input for the PCAs, we calculated genotype likelihoods for each of the three data sets using ANGSD (*73*), specifying the GATK algorithm (−GL 2) and the following parameters; only consider reads mapping to one location (−uniqueOnly 1), remove secondary alignments (− remove_bads 1), only include reads where both the forward and reverse reads mapped (− only_proper_pairs 1), a minimum mapping quality of 20 (−minMapQ 20), a minimum base quality of 20 (−minQ 20), a minimum minor allele frequency of 0.05 (−minmaf 0.05) skip triallelic sites (−skipTriallelic 1), remove SNP sites with a p-value larger than 1e^-6^ (−SNP_pval 1e-6), infer major and minor alleles based on genotype likelihoods (−doMaf 1), use the reference allele as the major allele (−doMajorMinor 4), output as beagle likelihood file (−doGlf 2), and specifying only sites found in at least 95% of total individuals (−minInd). Using the output beagle likelihood file, we computed the PCAs with PCAngsd (*74*) as well as the admixture proportions for the Greenland only dataset with the -admix parameter.

To quantify the divergence between individuals sampled from Kane Basin and Baffin Bay management units from west Greenland and between individuals sampled from the west and east coasts of Greenland, we used a consensus haploid base call in ANGSD and calculated the fixation index (*F*_ST_). We performed pseudo-haploid base calls to control for biases that may be introduced from the inclusion of the low-coverage data in this analysis. To compute this, we specified the following parameters in ANGSD; call the consensus haploid base (−dohaplocall 2), -uniqueOnly 1 -remove_bads 1 -only_proper_pairs 1 -minMapQ 20 -minQ 20 -GL 2 -minInd 50 -doMajorminor 1, only include an individual at a given site if the individual has a read depth of at least 3 (−setMinDepthInd 3). The resultant haploid output was run through the popgenWindows.py python script (https://github.com/simonhmartin/genomics_general), while specifying 200kb non-overlapping sliding windows in which to calculate *F*_ST_. We used a Wilcoxon signed rank test with continuity correction to test for significance from 0.

### Heterozygosity

Many of the individuals included in this study were of medium to low coverage (<20x). Therefore, to determine the number of individuals we could include in the heterozygosity analyses, we investigated the ability to accurately call genome-wide heterozygosity based on sequencing coverage. For this, we selected a single high-coverage individual (BGI-polarbear-PB_105, ∼30x), which we downsampled to 20x, 15x, 12x, 10x, 7.5x, 5x, 3x, 2x, and 1x using ATLAS (*75*). To account for variability in the subsampling, we repeated this step ten times, giving a total of 90 independent sub-samplings. Each of these subsamplings was then treated independently in the following analyses.

We inferred recalibration parameters to correct for sequencing errors on each of the 90 sub-samplings in ATLAS (task=recal), only specifying the mitochondrial genome, a minimum base quality of 20 (minQual=20), and equal base frequencies (equalBaseFreq). We only selected the mitochondrial genome as recal uses haploid sites as a training set to infer the recalibration parameters, which can then be used to correct for errors at a genome-wide scale. We estimated heterozygosity for each downsampling in ATLAS (task=estimateTheta) (*76*) using the recalibration parameters previously calculated, a minimum base quality of 20, and only including the first 80 Mb from each pseudo-chromosome from pseudo-chromosomes 1-18 (out of 21 available) (*21*). We plotted the resultant proportion of “true” heterozygosity recovered at the different downsamplings, assuming 20x was representative of the true heterozygosity. We used a polynomial regression line with an order of two (Supplementary Fig. S10) to calculate an equation to correct for false negative heterozygosity caused by decreasing coverage. Due to the high variation in heterozygosity at 1x and 2x, we excluded these sub-samplings from this calculation. From this, we used a correction for false negative heterozygosity in samples <20x using the equation (−0.0004 x coverage^2^) + (0.0218 x coverage) + 0.7075. We note that this equation may be very software, parameter, and data specific, and should be implemented with caution on other datasets and software.

Based on the variability around estimates when downsampling to 3x, we only calculated the heterozygosity for every individual with >5x average coverage (n = 20) in ATLAS, using the same parameters as above. Furthermore, to make the results as comparable as possible, we downsampled any individual with >20x coverage to 20x. The heterozygosity of any individual <20x was corrected using the above equation. The 95% confidence intervals around the heterozygosity estimates were taken from the Fisher confidence intervals given by ATLAS.

### Nucleotide diversity

The popgenWindows.py python script used for the *F*_ST_calculation above, also outputs nucleotide diversity. We used a Welch unpaired two sample t-test to test for significant differences in nucleotide diversity estimates between the individuals sampled in west Greenland and those sampled in east Greenland in R v4.1.1 (*77*).

### Inbreeding

To estimate the inbreeding coefficients of each individual Greenlandic bear, we used NGSrelate v2 (*78*) and genotype likelihoods as input, which we computed using ANGSD. As NGSrelate v2 has been shown to give accurate results from individuals down to 1x, we used our complete Greenlandic bear dataset, numbering a total of 85 individuals. To compute the genotype likelihoods, we used the same ANGSD parameters as the PCA, including sites where at least 1 individual had coverage (minInd 1), and output the genotype likelihoods as binary (−doglf 3) as required for NGSrelate.

### Demographic reconstruction

To reconstruct deep-time demographic trajectories, we implemented a Pairwise Sequentially Markovian Coalescent (PSMC) model (*33*). To create the input file for the PSMC analysis, we used ATLAS. PSMC relies on the distribution of heterozygosity across the genome of a single individual, and therefore requires heterozygous sites to be called as accurately as possible. Because of this requirement, we only performed PSMC on the individuals that had >20x. For comparability we selected six individuals from each management unit (Kane Basin n = 6; Baffin Bay n = 6, east Greenland n =6). Furthermore, to ensure sample comparability, we downsampled all individuals >20x to 20x coverage using ATLAS. We computed the input psmcfa file using recalibration parameters based on the mitochondrial genome, a minimum base quality of 20, and only including the first 80 Mb of each pseudo-chromosome from pseudo-chromosomes 1-18 (out of 21 available) as done previously to avoid assembly problems (*21*). The output psmcfa file was run through PSMC specifying atomic intervals 4+25*2+4+6, and plotted using a generation time of 11.2 years and a mutation rate of 1.83×10^−08^ per generation (*21*).

To estimate the more recent demographic histories of bears sampled from the west coast and east coast of Greenland over the last 20,000 years, we implemented independent stairway plots (west *vs* east) based on the unfolded site frequency spectrum (SFS) specifying the spectacled bear as the ancestral state and only using individuals with >20x coverage. We repeated the analysis using the giant panda as the ancestral state to assess the impact of the species used to designate the state. We calculated the SFS for each coast independently from allele frequencies on a representation of the whole genome (pseudo-chromosomes 1-3) in ANGSD (−doSaf 1), and specifying the following parameters: only include sites covered in all individuals (−minind), the ancestral sequence as the giant panda (−anc), -uniqueOnly 1, - remove_bads 1, -only_proper_pairs 1, -minMapQ 20, -minQ 20, -doMajorMinor 4, -doMaf 1, - skipTriallelic 1, -GL 2, -doGlf 2, -SNP_pval 1e-6. We used winsfs to convert the ANGSD allele frequencies into the SFS (*79*). We plotted the results using a generation time of 11.2 years and a mutation rate of 1.83×10^−08^ per generation (*21*).

The ancestral sequence was specified using the -anc parameter in ANGSD, and using a fasta file of either the spectacled bear or the giant panda. To obtain the fasta files, we mapped the spectacled bear (Genbank Bioproject: PRJNA472085) (*80*) and giant panda raw reads (Genbank Bioproject: PRJNA38683) (*81*) to the polar bear genome using BWA and used ANGSD to generate a consensus base call fasta file (−doFasta 2) with the filters: -mininddepth 5 -minq 25 - minmapq 25. We further tested the robustness of the stairway plot results based on the number of samples by downsampling the number of individuals from west Greenland to nine individuals, and rerunning the same analysis.

To corroborate with the stairway plot demographic results, we calculated Tajima’s D for each population in ANGSD (*82*). We limited our dataset to only individuals with >3x coverage and calculated the allele frequencies and SFS for each population as done above and with the spectacled bear as the ancestral state. To control for variation across the genome we calculated Tajima’s D in 10 Mb windows and extracted the mean values. Significance from 0 was calculated using a Wilcoxon signed rank test with continuity correction. A positive Tajima’s D indicates a recent demographic contraction, whereas a negative value indicates a demographic expansion (*35*).

### Stable isotope analyses

Approximately 100 mg of powdered bone was removed from each polar bear skull, as well as the pinniped skulls. The powdered samples were sonicated in a 2:1 chloroform methanol (v/v) solution for 1 h to remove lipids (*83*). The solvent was removed and the samples were dried under normal atmosphere for 24 h. The bone was then demineralized in 0.5 M HCl for 4 h under constant motion (orbital shaker), rinsed with Type I water (18.2 MΩ·cm), and then heated at 75°C for 36 h in 3.5 mL 0.01 M HCl to solubilize the collagen. The solution containing the collagen was freeze dried and then suspended in 1.6 mL of 75°C Type I water. This solution was again lipid extracted by adding 2 mL methanol and 4 mL chloroform (2:1:0.8 chloroform:methanol:water, (*84*) and sonicating for 1 h in a culture tube (Tube I). The samples were centrifuged and the top layer containing the collagen was collected and transferred to a new culture tube (Tube II). 4 mL of Type I water was added to Tube I to recover any residual collagen, centrifuged, and the top layer was again added to Tube II. Tube II was heated at 60°C for 4 h to remove methanol from the solution, and the remaining solution was then freeze dried.

Carbon and nitrogen isotopic and elemental compositions were determined using a Thermo Delta V isotope ratio mass spectrometer (IRMS) coupled to a Costech 4010 elemental analyzer (EA) or a Nu Horizon IRMS coupled to a EuroEA 3000 EA. In both cases *δ*^13^C and *δ*^15^N values were calibrated relative to the international reference scales (VPDB and AIR) using USGS40 and USGS41a (*85, 86*). Measurement uncertainty was assessed using three in-house standards with the following established isotopic compositions: SRM-1 (caribou bone collagen, *δ*^13^C = -19.36±0.11 ‰, *δ*^15^N = +1.81±0.11 ‰), SRM-2 (walrus bone collagen, *δ*^13^C = - 14.77±0.11 ‰, *δ*^15^N = +15.59±0.11 ‰), SRM-14 (polar bear bone collagen, *δ*^13^C = -13.67±0.07 ‰, *δ*^15^N = +21.60±0.15 ‰), and SRM-15 (phenylalanine, *δ*^13^C = -12.44±0.04 ‰, *δ*^15^N = +3.08±0.12 ‰). Twenty percent of the samples were analysed in duplicate. Standard uncertainty was calculated to be ±0.14 ‰for *δ*^13^C and ±0.28 ‰for *δ*^15^N (*87*).

The *δ*^13^C value of atmospheric CO_2_ has declined significantly since the 1800s due to the combustion of isotopically light fossil fuels (*88*), a process known as the ‘Suess Effect’. Accordingly, when comparing *δ*^13^C data derived from fauna sampled across recent decades or centuries it is necessary to perform a correction to account for this change. The isotopic composition of dissolved inorganic carbon reservoirs in the ocean takes time to equilibrate with that of the atmosphere and this varies among different ocean basins (*89*). We adjusted the *δ*^13^C values of all of the samples (polar bears and prey) according to the following equation (*90*):

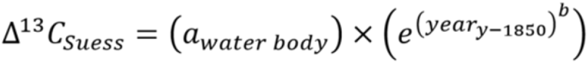

*Δ*^13^C_*suess*_ is the difference in *δ*^13^C between the year in question and 1850 where *b* is 0.027 ‰, fitting the curve to the decline in global oceanic *δ*^13^C (*91*) and *a*_*water body*_ is the annual rate of *δ*^13^C decrease, in this case set to 0.014 ‰for the North Atlantic (*92*). This correction was applied to all polar bear and prey species. Where the precise year of collection was uncertain (e.g., 1928-1929) we used the midpoint of the range given. Since only older samples had these uncertain collection dates and the magnitude of the Suess Effect correction is small for these samples, a variance of a few years was inconsequential.

We assessed whether we could pool individuals from the two management units (Kane Basin and Baffin Bay) from west Greenland by comparing *δ*^13^C and *δ*^15^N using Mann–Whitney– Wilcoxon and Student’s t-tests depending on whether the data satisfied normality and homogeneity of variances with the statistical package R (*77*). Bone collagen *δ*^13^C and *δ*^15^N did not differ significantly between Kane Basin and Baffin Bay (W = 27, P = 0.45 for *δ*^13^C, and t = - 1.62, df = 11, P = 0.13 for *δ*^15^N). Therefore, all individuals sampled from west Greenland were pooled together for subsequent analyses. Movement of individuals between the regions has been shown via telemetry (*93*), further supporting this decision.

To assess differences in polar bear isotopic compositions according to sex and geographic location (west *vs* east), we used either a Mann-Whitney U test (if distributions were not normal as determined by a Shapiro-Wilk test), a Student’s t test (if distributions were normal and variances were equal), or a Welch’s t test (if distributions were normal and variances were unequal as determined by an F test).

To determine the relative contributions of marine mammal prey species to the diets of the bears, we used a Bayesian mixing model (MixSIAR) (*94*) with uninformative priors. Because bone collagen of predator and prey were being compared, trophic discrimination factors +0.50±0.25 ‰for *Δ*^13^C_predator collagen-prey collagen_ and +4.0±0.5 ‰*Δ*^15^N_predator collagen-prey collagen_ (*95*).

### Geometric morphometric analyses

We defined 32 cranial landmarks that could be unequivocally located and were presumed to be homologous among all skulls (Supplementary Fig. S11 and Supplementary text). Three-dimensional coordinates of the landmarks were registered with a Microscribe® 3D digitizer on all skulls by the same observer. The raw landmark coordinates were run through the generalized least-squares Procrustes superimposition (Rohlf & Slice, 1990) using the MorphoJ-program (*96*), which was also used for analysis of geographical shape differences. The Procrustes procedure used here was amended by the suggestions of Klingenberg *et al*. (*97*), to deal with the redundancy of data points caused by the object symmetry of the vertebrate skull. To exclude size-related variation, all further analyses on shape were performed on the residuals of a multivariate regression of shape (Procrustes coordinates) on the logarithm of centroid size (the square root of the summed squared distances of each landmark to the averaged coordinates of the configuration).

To ensure independence of vectors describing sexual and geographical (west *vs* east) differences before pooling sexes, the directionality of the vector describing shape differences between sexes in the larger sample from east Greenland was compared to the vector describing shape differences between individuals from west and east Greenland (both vectors defined by linear discriminant analysis). If these vectors were unrelated, sex-related and geographic differences are unrelated and could hence be analysed independently with sexes pooled. The vectors describing the shape differences between west and east Greenland individuals, and between east Greenland males and females (where the sample size allowed us to test for differences between sexes), were largely independent at 87.0° (with 90° representing complete independence). We were therefore able to pool males and females in comparisons of shape differences between individuals to increase sample sizes.

As a previous study (*98*) showed no obvious morphological differences between bears found at different localities along the east Greenland coast, we pooled all east Greenland individuals together. Before comparing east and west Greenland individuals, we tested for differences among the west Greenland individuals, as Kane Basin and Baffin Bay bears are regarded as different management units. There were 14 bears from each of these areas, and a permutation test for differences gave a p value of 1. The Procrustes distance between management unit mean shapes was 0.012 as compared to a distance between all west Greenland and east Greenland individuals of 0.036. Based on this, west Greenland bears were pooled for further analyses.

Shape differences between west and east Greenland were investigated by a linear discriminant analysis with Mahalanobis distance based reclassification of each specimen to the most likely area by jackknife cross-validation (*99*). A permutation test with 1,000 runs was used to assess the statistical significance of shape differences between the two geographical (west *vs* east) samples. Furthermore, using the statistical package R (*77*), Student’s *t*-tests were used to compare centroid sizes between both sexes from the two areas. The variation of skull shapes within each area was assessed by computing the Procrustes distance of each specimen to the respective group mean shapes. As for centroid sizes, Student’s *t*-tests were used to compare the levels of individual variation of skull shape between regions.

### Habitat suitability modelling

#### Occurrence records

Occurrence records were compiled for the polar bear from the early 20^th^ century to the end of 2019. All records were obtained from publicly available databases (e.g. GBIF) or digitised from the literature (Supplementary Table S5). In some cases figures were georeferenced and points extracted manually (e.g. Laidre, *et al. (93)*). If the density of points was too large for manual extraction (e.g. Durner, *et al*. (*100*)), polygons bordering the points were digitised, with random points placed inside the boundary of the polygons. The number of points within each polygon was maximised using a criterion that all random points had to be separated by a minimum distance of 100 km. Collation of records from all sources provided in Supplementary Table S5 resulted in 139,641 occurrences between 1910 and 2019.

Occurrence records were then thinned spatially and temporally to reduce the number of samples. Records before 1985 and after 2014 were excluded and occurrence records that occurred on land or outside the boundary of the 19 recognised polar bear management units (*25*) were removed. We spatially thinned the remaining records (n = 1,724). This was done using the *spThin* package for R (*101*). A minimum distance of 100 km between records was permitted, with the process repeated 100 times. Spatial thinning resulted in *n* = 583 records covering the circumpolar region (Supplementary Fig. S12).

#### Climate data

Climate data for the analysis came from two separate sources which permitted matching the contemporary occurrence records to contemporary climate data whilst also permitting hindcasts of habitat suitability to be generated.

Contemporary climate was taken from historical simulations from the Hadley Centre HadGEM3-GC3.1 model (*102*) produced for the Coupled Model Intercomparison Project Phase 6 (CMIP6) (*103*). The historical simulations done using the HadGEM3-GC3.1 model have good agreement with observed climate trends and variability for sea-surface temperature and Arctic sea-ice over the 1850-2014 simulation period (*103*). Data on sea-surface temperature (SST), sea-surface salinity (SAL), and sea-ice concentration (SIC) were downloaded from the Earth System Grid Federation CMIP6 data portal (https://esgf-node.llnl.gov/search/cmip6/) on the models native grid, before being re-gridded to a global 1° x 1° regular grid using bilinear interpolation.

Paleoclimate data were accessed using a 1° x 1° oceanic climate dataset for the period 11 thousand years ago (kya) to present (1950 C.E.) following Armstrong, Hopcroft and Valdes (*46*). These data were generated by temporally linking 42 discrete snapshot simulations from the HadCM3B-M2.1 coupled general circulation model (*104*). The HadCM3B-M2.1 model has been shown to accurately represent different aspects of the climate system including sea surface temperatures and ocean circulation (*104*). The snapshot simulations were linked using splines based on monthly climatologies, before interannual and millennial scale variability (e.g. Dansgaard-Oeschger (*105*) and Heinrich (*106*) events) were imposed on the timeseries. The data was then bilinearly downscaled to 1° x 1° with SST and SAL data bias-corrected using a simple delta (change-factor) method following Armstrong, Hopcroft and Valdes (*46*) with simulated data being harmonised against the World Ocean Atlas 2018 dataset (*107*). SIC was harmonised against the 20^th^ Century Reanalysis Dataset (*108*) using a climatologic period of 1850-1950 C.E. Following Armstrong, Hopcroft and Valdes (*46*) the multiplicative bias correction was capped at 3x the simulated value.

To harmonise the two datasets, permitting seamless hindcasts of habitat suitability, the CMIP6 data was harmonised against the bias-corrected paleoclimate data. This was done using the same simple delta method used by Armstrong, Hopcroft and Valdes (*46*) and a climatological period of 1850-1950 C.E. SST data was corrected using an additive approach, with salinity and sea-ice fraction corrected using a multiplicative approach due to both variables being bounded by 0. Corrected sea-ice cover values that exceeded 100% were then truncated back to 100%. The resulting data was a continuous time series of maps of climatological averages, calculated over a 30-yr window, with a step of 50 years, for the period 11 kya to 0 kya. Climatological averages for SST, SAL, SIC, as well as two bioclimatic variables (*109*) were then calculated using the harmonised data (Supplementary Fig. S13) between 1985 and 2014. We calculated SST seasonality (BIO 4) as the standard deviation of monthly temperatures (x 100), and then calculated the coefficient of variation in annual sea-ice cover (similar to BIO 15; (*109*)). All contemporary occurrence records were then matched to this data and used to calibrate the ecological niche model.

We opted to use SST, SAL, SIC, and the two bioclimatic variables (BIO 4, CV sea-ice cover) in our analysis, as we were limited with regards to the type of information we could calculate for our hindcasts. As such, we were limited to using more distal predictors which are typical in species distribution models instead of proximal predictors (i.e. predictors that directly influence distributions and demographic processes) (*110*). In extreme environments where species are not occupying their optimal realised niche, distal variables are theorised to be as successful as more direct, proximal variables in predicting the relationships between environmental pattern and process (*111*), with these variables acting as proxies for proximal constraints such as prey availability (*112*). Because robust spatiotemporal data were not available to generate some proximal predictors of polar bear geographic range (e.g., prey abundance, accurate hindcasts of bathymetric data), we used available distal surrogates. While previous work has shown that polar bears have a preference for habitats with ≥ 50 % sea-ice cover, particularly over the continental shelf (*100*), we were not able to calculate these metrics (e.g. bathymetric depth, distance from continental shelf) as we did not have access to accurate bathymetric data for our paleoclimate reconstructions (*46*). SST has previously been linked to higher δ^15^N in ringed seals affecting prey quality (*113*), is known to affect seasonal swimming behaviour of polar bears (*114*), with warmer SST impacting the timing of sea-ice formation (*115*). Finally, salinity has been linked to regional differences in ocean productivity with decreased diversity and biomass of the pelagic and benthic prey taxa of Arctic marine mammals in areas of lower salinity (*116*). Consequently, our estimates of SST, SAL, SIC, BIO 4, and CV sea-ice cover metrics could be considered both proximal and distal predictors as they have a direct (proximal) influence on polar bear physiology and behaviour (and therefore fitness), and an indirect (distal) influence on prey distributions.

#### Ecological niche model

We created an ecological niche model (ENM) for the polar bear using the Maxnet package for R (*117*). The correlative ENM approach used here assumes that species are in equilibrium with the environment, and thus their observed distribution is a good indicator of their ecological requirements (*112*). Maxnet creates maximum entropy models (hereafter MaxEnt) using penalised generalised linear models (*118*). We tuned and did cross-validation of our MaxEnt model using the ENMeval package for R (*119*).

##### MaxEnt model

MaxEnt models require background points. We set the number of background points to 10,000 random locations constrained to an area defined as the minimum bounding polygon of the polar bear subpopulation management units (*25*). Due to the autocorrelated nature of ecological data, spatially explicit cross-validation folds are necessary to assess the transferability of ecological niche models to new locations and climates. We defined our 5 cross-validation folds using the blockCV package for R (*120*). We opted to use 5 blocks as there are 5 sea-ice ecoregions in the circumpolar zone (*121*). Blocks were assigned randomly using a distance of 1,000 km. We also performed 5-fold cross validation on the spatial blocks to achieve the most even spread of occurrence and background points within each of the 5 blocks. Occurrence and backgrounds points were subsequently assigned to spatial blocks which were used as holdout test sets during model cross-validation – e.g. the model was built using data from blocks 1-4 and tested on 5, the next model was built on blocks 2-5 and tested on 1, etc.

MaxEnt is capable of building complicated nonlinear response curves to environmental data using a variety of feature classes (i.e. functions and transformations of the data (FC) and penalisation values (*37*). To reduce overfitting, a regularisation parameter (RM) that attempts to balance model fit against complexity was done by penalising models based on the magnitude of their coefficients. Six different FCs are available in MaxEnt models: linear (untransformed), quadratic (square of the variable), product (product of two variables – analogous to an interaction term in regression), hinge (a “step” function with a linear function above a critical threshold), threshold (a “step” function generating a constant function above a critical threshold), and categorical (*37, 118, 122*). We used several FC combinations and RMs from 1 to 5 in 0.5 increments. The FC combinations were: linear only (L); quadratic only (Q); hinge only (H); linear and quadratic (LQ); quadratic and hinge (QH); linear, quadratic, and product (LQP); quadratic, hinge, and product (QHP); and linear, quadratic, hinge, product (LQHP; the MaxEnt default). Correlations between variables were checked following Dorman et al. (*123*), however MaxEnt is able to handle correlated predictors, and through penalisation (RM), can directly account for redundant variables (*124*).We selected for the optimal combination of RM and FC by maximising model spatial transferability (i.e. low AUCdiff scores; (*125*)), and minimising the 10% omission rate (OR10) whilst maximising AUC_test_ (*126*). However, to break ties in our results we further minimised the standard deviation in OR10 (from the cross-validation test/train) and maximised the RM. This process ensured we had the most heavily penalised, but most transferable model.

To restrict extrapolation of the ENM to novel environments, all models were clamped during both model building and spatial projection. Clamping forces the response of the variables to be at a constant value at the limits of the training range – effectively holding species responses at constant probabilities (*122*). Clamping is particularly important during hindcasting and forward-projecting when it is unlikely that the fundamental niche (or a close enough approximation) has been captured (*127*) – which is unlikely using contemporary occurrence data only.

Following model building and selection, spatial projections of contemporary and past suitability were produced using the appropriate data sources. Projections were then rescaled from the default exponential output to 0 and 1. To identify upper and lower boundaries for the rescaling we calculated the OR10 and OR90 using the contemporary occurrences. The OR10 is defined as the threshold which excludes regions with suitability values lower than the values for 10% of occurrence records – the assumption being that the lowest 10% of occurrence records are from regions that are not representative of the species overall habitat and can be omitted. The OR90 is the opposite – the threshold above which the upper 10% of occurrence records occur.

The assumption here is that the upper 10% of occurrence records are from highly suitable habitat and can thus be considered the maximum value. Projections of climate suitability were then scaled between 0 and 1 following the equation:

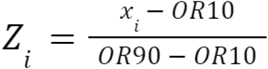

Where *x*_*i*_ = climate suitability at grid cell *i*, OR10 and OR90 are as defined above. To ensure that hindcasts of suitability were relevant to the species contemporary habitat suitability values, the OR10 and OR90 thresholds were calculated on the contemporary projection and applied to all hindcast projections.

Following the creation of our hindcasts we analysed the centre of gravity for each generational step of our habitat suitability models for the Greenland region. The centre of gravity was calculated as the weighted latitude of the habitat suitability projections using the following equation:

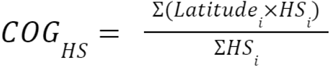

Where COG_HS_ is the weighted centre of gravity, and Latitude_i_ and HS_i_ are the latitude and habitat suitability at grid cell *i*.

## Acknowledgements

We would like to acknowledge the large number of local hunters as well as Jan Lorentzen and the late Jonas Brønlund who conducted the polar bear tissue sampling, and organised the collection and shipment of samples in east Greenland. We thank Lars Johannessen for assistance with seal samples from the National History museum, University of Oslo (loan number: DMA-30). Finally we would like to thank Daniel Klingberg Johansson for assisting in locating the pinniped skulls at the Natural History Museum of Denmark.

## Author contributions

Conceptualization, MVW, EDL; Formal analysis, MVW, SCB, SO, MBS, SCB, SON, JASC, AG, PS; Investigation, MVW, JL, MBS, JM, SKB, ML, AAC; Writing – Original Draft, MVW, SR, AG, EDL; Writing – Review & Editing, All authors; Funding Acquisition, EDL; Resources, EA, RD, CS, PS, EDL; Supervision, DF, PS, EDL.

