## Supplementary information for "Impact of Holocene environmental change on the evolutionary ecology of an Arctic top predator"

**Supplementary table S1:**
Sample overview and mapping statistics of the 107 Atlantic Arctic polar bear individuals analysed in the study, and of the giant panda used to infer the ancestral state in the stairway plot analysis. Locality information is provided, including the putative management unit of the individual samples.

**Supplementary table S2:**
Corrected autosome-wide heterozygosity of the 24 polar bear individuals with >5x coverage.

**Supplementary table S3:**
Voucher specimen information and stable isotope values of the 31 polar bears and 110 pinniped specimens, representing five polar bear prey species, analysed in this study. '$\delta^{13}$C Suess' represents the Suess-corrected $\delta^{13}$C values. All specimens are either housed at the Natural History Museum of Denmark (NHMD), University of Copenhagen or the National History museum (NHMO), University of Oslo. The management unit of each individual polar bear sample was determined based on the sampling locality, where known.

**Supplementary table S4:**
Voucher specimen information of the 134 polar bear skulls used in the geometric morphometric analyses. Blank data points represent unknown values. All samples are housed at the Natural History Museum of Denmark (NHMD), University of Copenhagen. As soft tissue specimens exist at Aarhus University (AU) for some of the polar bear specimens, field number is included, which corresponds to the unique identifier used by AU for each polar bear individual. Polar bear management unit was determined based on the sampling locality of the individual. If exact locations were not available, the management unit was taken as the most probable based on rough geography. The management unit of each individual polar bear sample was determined based on the sampling locality, where known.

**Supplementary table S5:** Sources for the polar bear occurrence records used for ecological niche modelling.

**Supplementary figures**

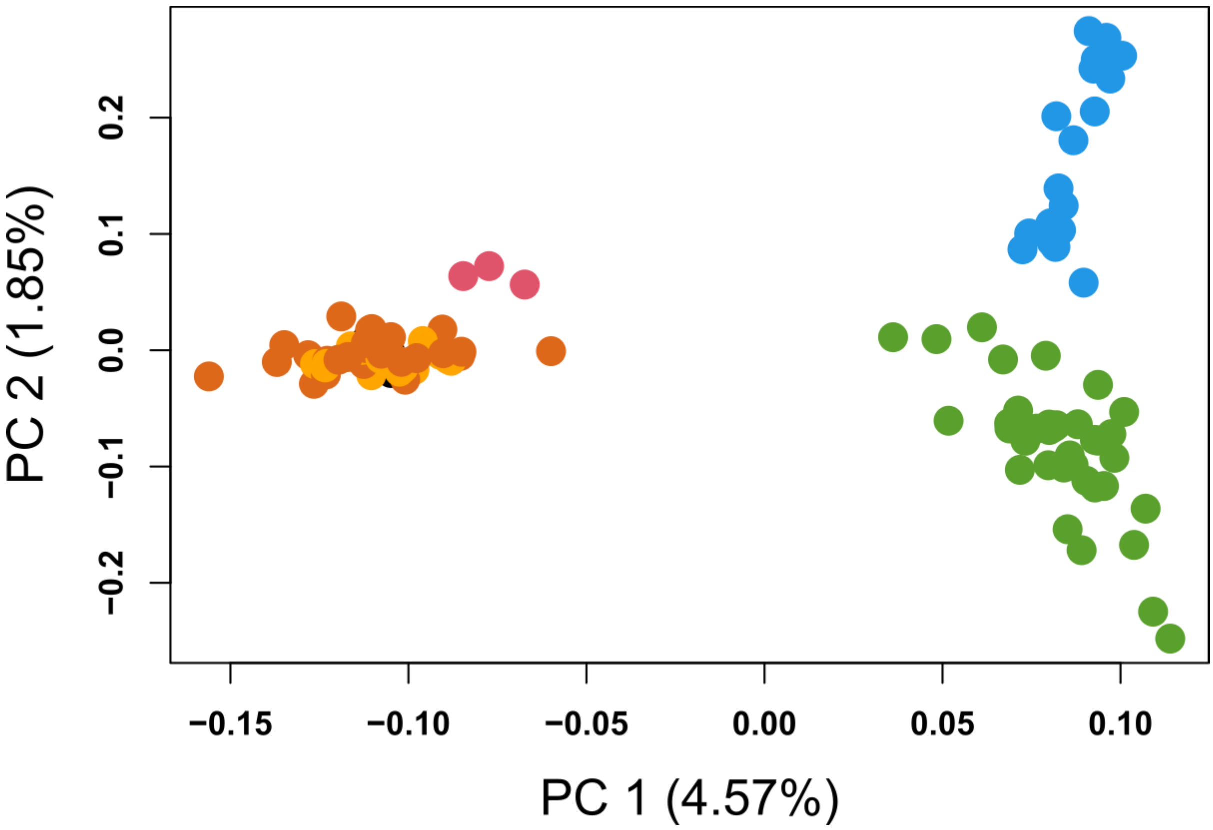


**Supplementary figure S1:** Principal component analysis using all 101 polar bear genomes from Greenland and surrounding areas, with the percentage of variance explained by each component indicated in parentheses. West Greenland (Kane Basin, dark orange, n = 27 ; Baffin Bay, light orange, n = 18; Unknown, black, n = 3), east Greenland (green, n = 37), Canada (red, n = 3), Svalbard (blue, n = 18).

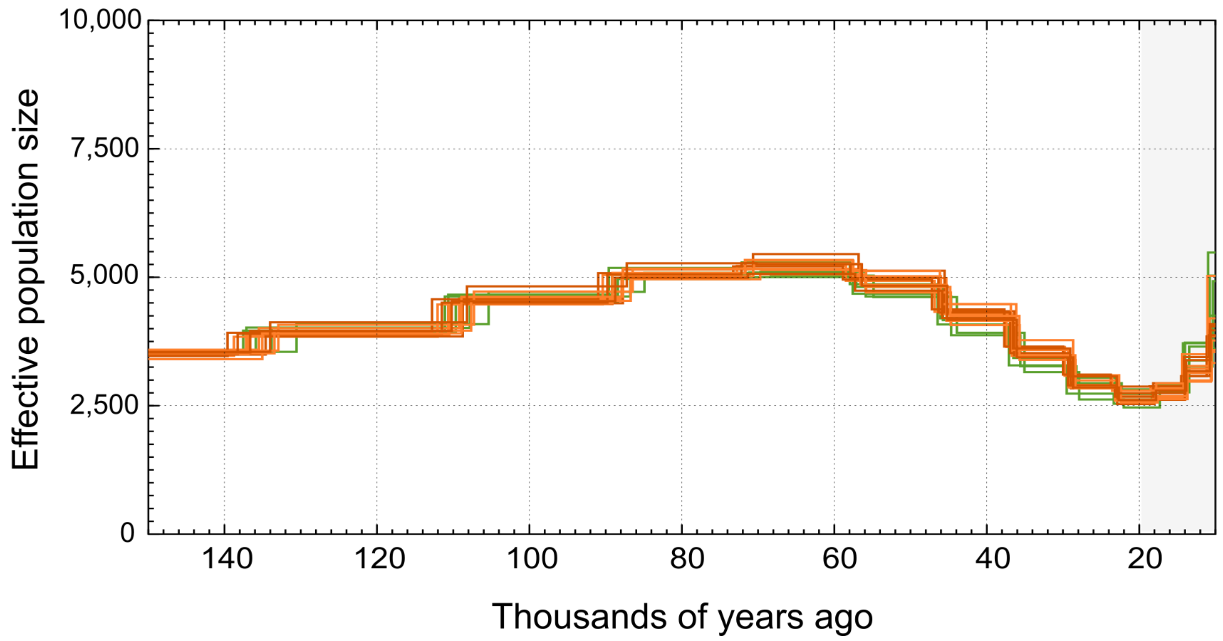


**Supplementary figure S2:** Changes in effective population size ($N_e$) through time inferred using PSMC on six individuals per management unit with genome-wide coverages >20x. The most recent ten thousand years are not shown due to the inaccuracy of this method for more recent time periods. West Greenland (Kane Basin, dark orange; Baffin Bay, light orange), east Greenland (green).

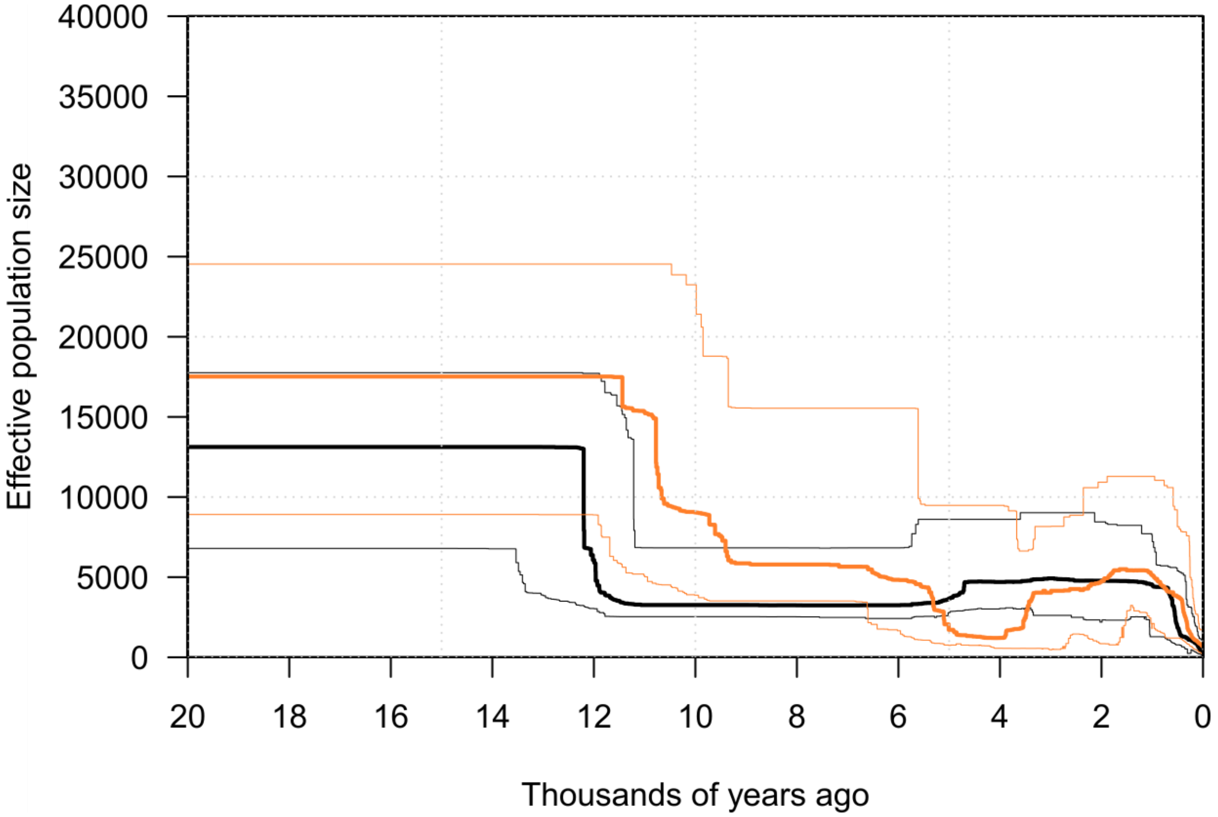


**Supplementary figure S3:** Demographic history of polar bears sampled from the west coast of Greenland calculated using stairway plots and the complete dataset of twelve individuals (orange), or a subset of nine individuals (black).

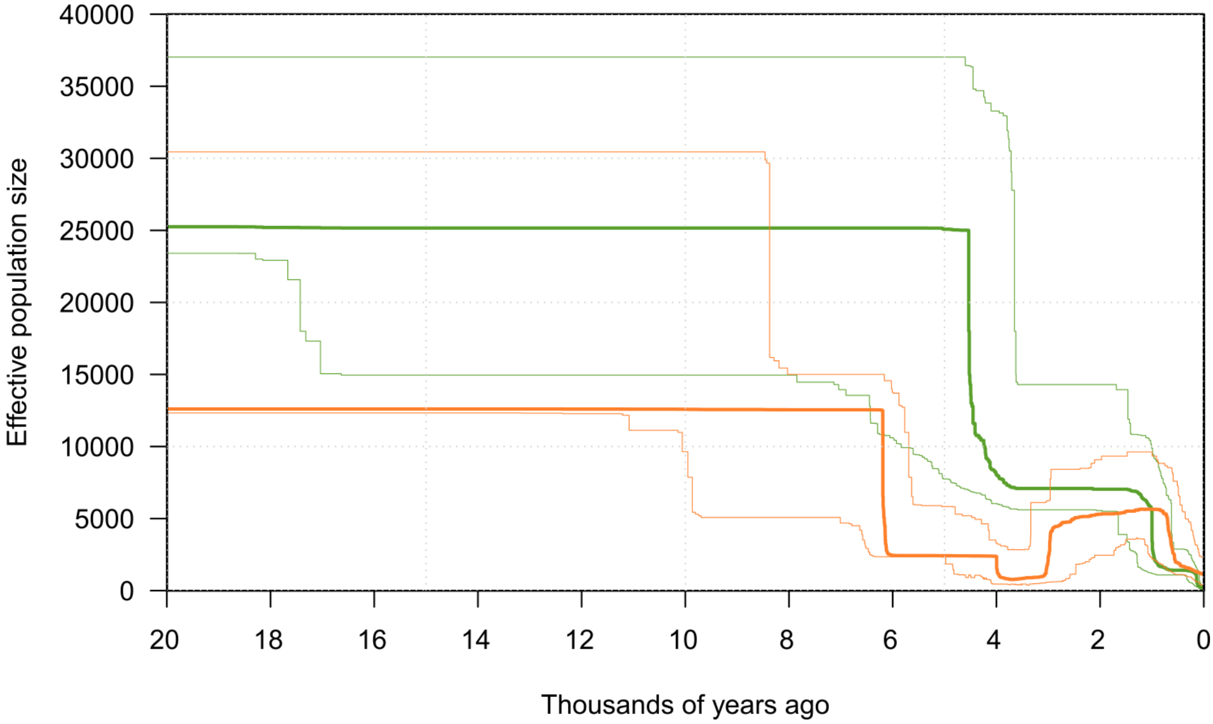


**Supplementary figure S4:** Demographic history of polar bears sampled from the west coast (orange) and the east coast (green) of Greenland calculated using stairway plots with the giant panda specified as the ancestral state when calculating the site frequency spectrum.

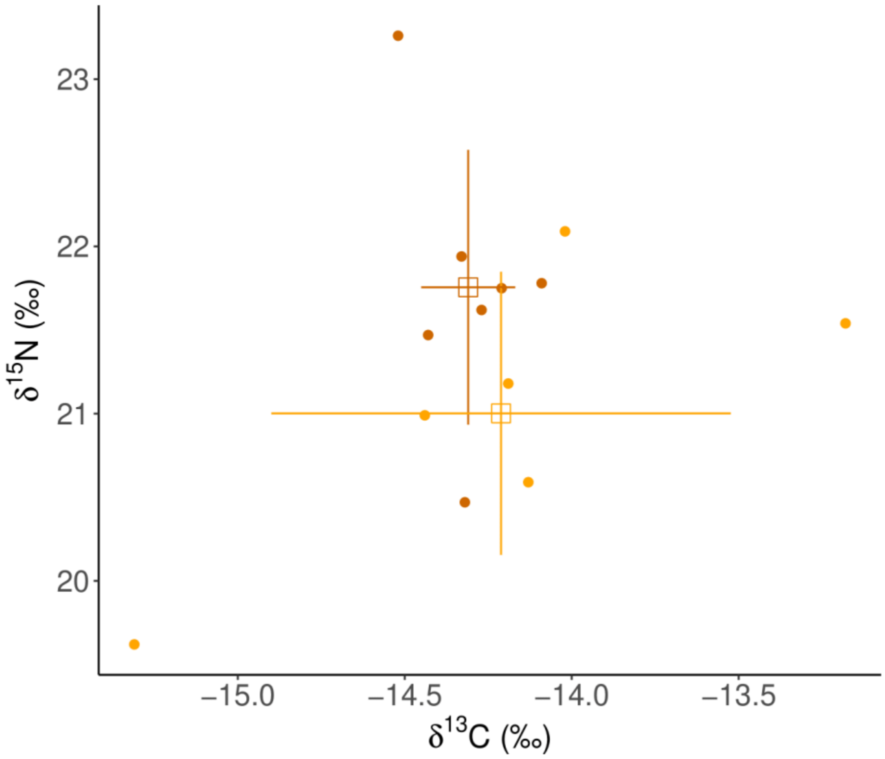


**Supplementary figure S5:** Stable isotopes for polar bears sampled from the west coast of Greenland. Colours indicate the management unit of the bear, dark orange is Kane Basin and light orange is Baffin Bay.

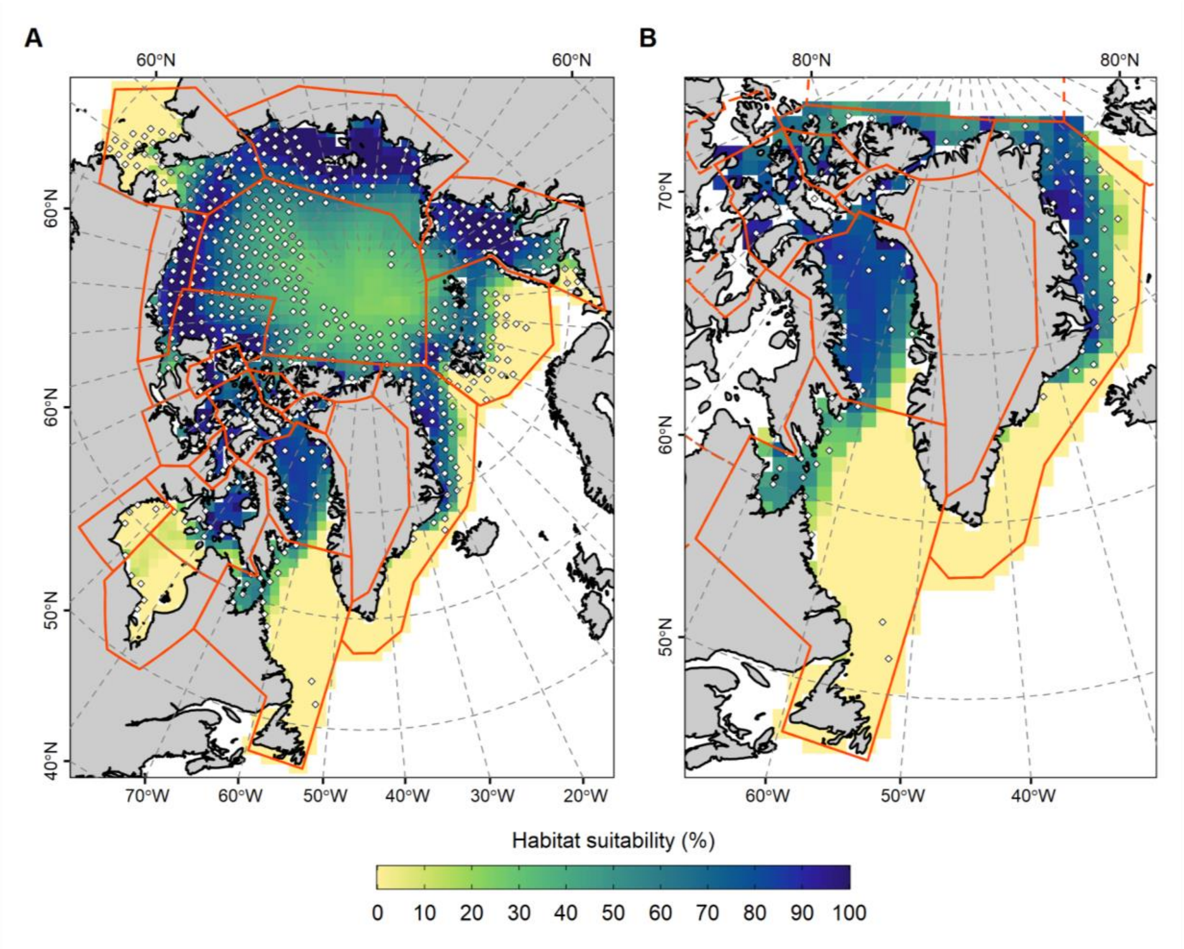


**Supplementary figure S6:** MaxEnt projections of habitat suitability for the polar bear based on contemporary occurrences and climate data. (A) Circumpolar habitat suitability values with occurrence records shown by white points, polar bear subpopulation management units are outlined in orange. (B) Habitat suitability values masked to the region surrounding Greenland.

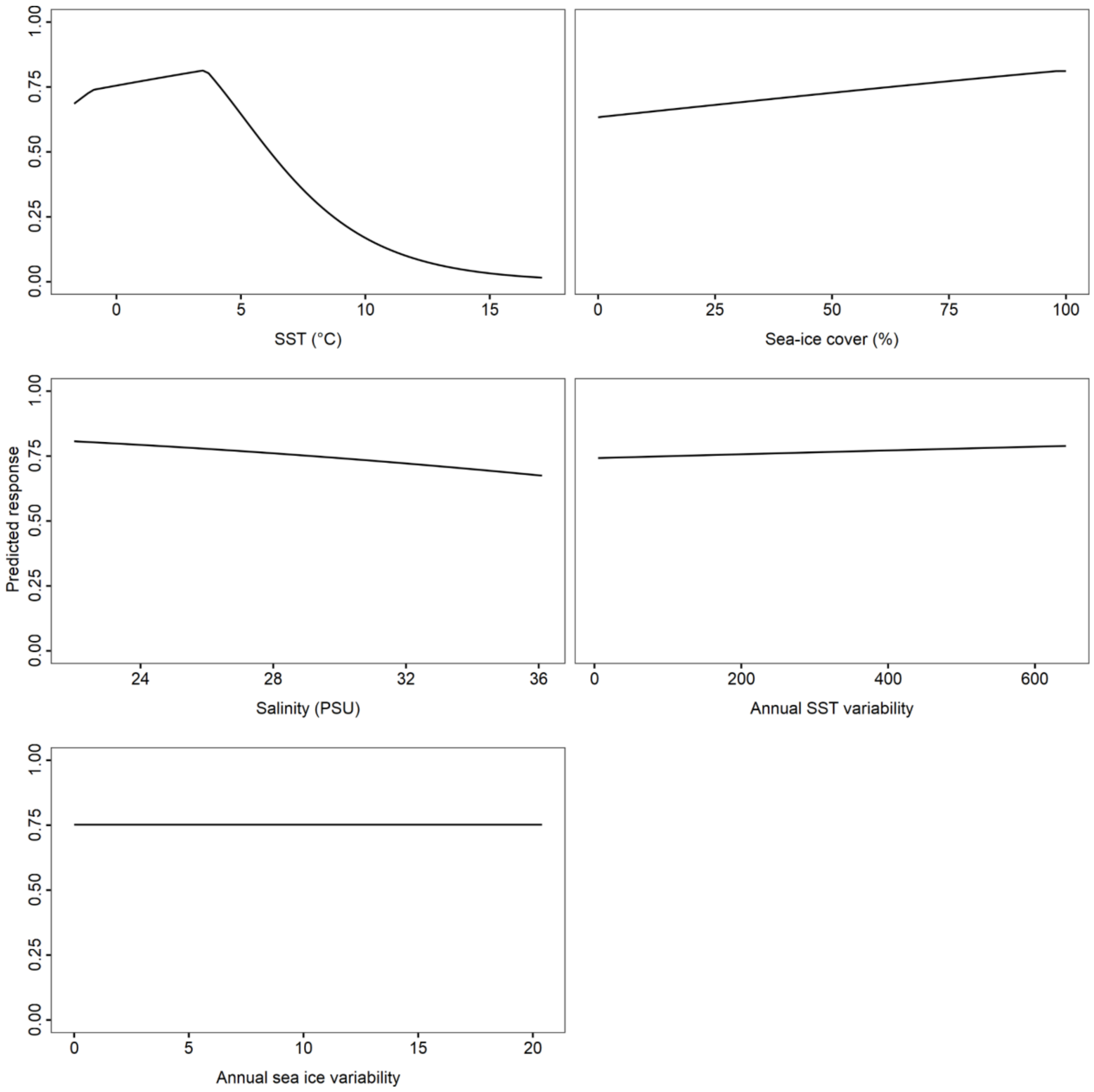


**Supplementary figure S7:** Response curves illustrating the relationship between MaxEnt predicted probability of occurrence and environmental variables. The values shown on the y-axis of the plots show the predicted probability of polar bear occurrence, for a model built using only that particular predictor variable.

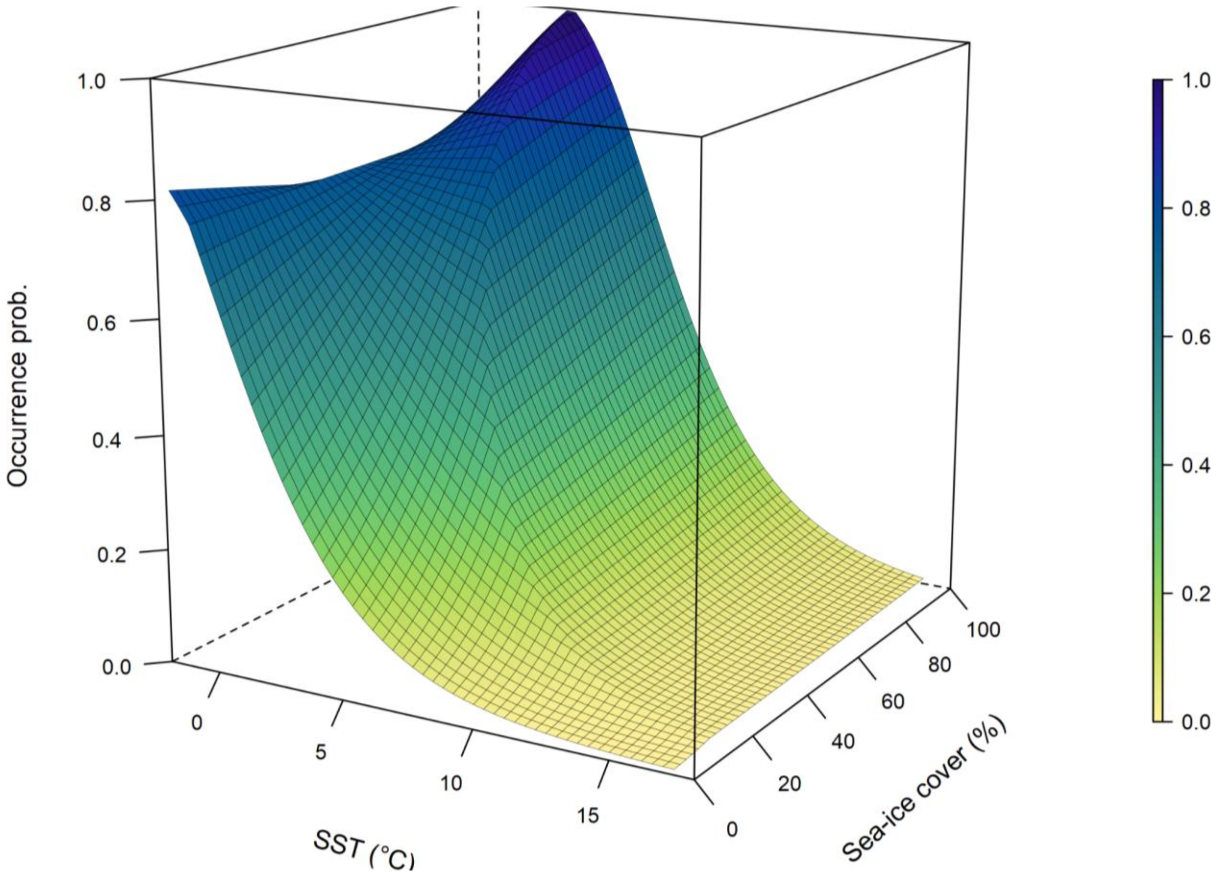


**Supplementary figure S8:** Probability of occurrence as a function of sea surface temperature (SST) and fractional sea-ice cover. The plot shows the marginal response of occurrence probability based on variation in SST and ice cover. Other predictors (salinity, SST seasonality, and coefficient of variation sea-ice cover) are held at their mean values.

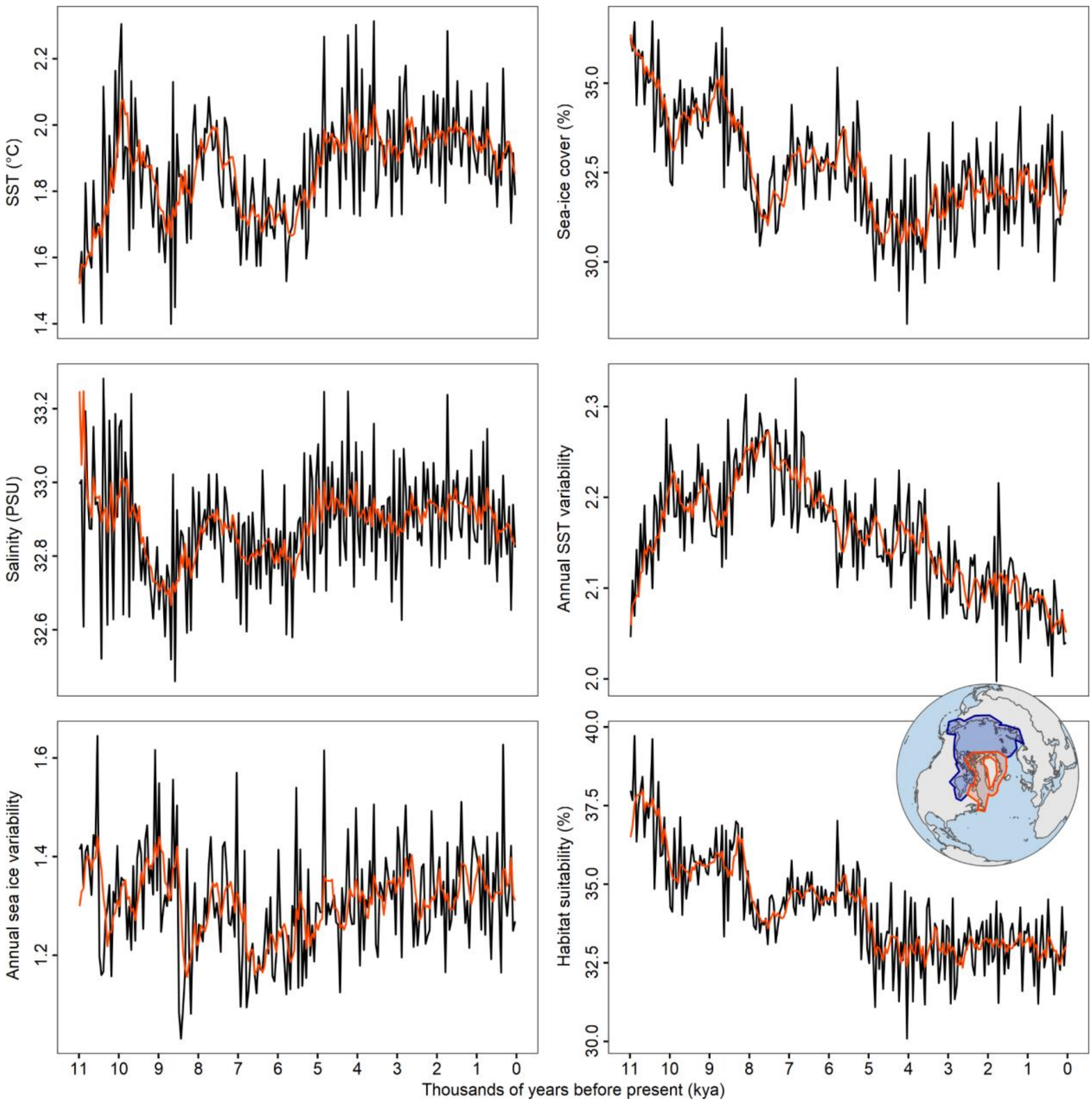


**Supplementary figure S9:** Areal means of the variables used in the model and predicted habitat suitability from 11 to 0 thousand years before present. The panels show the areal mean values in sea surface temperature (SST), sea-ice cover, salinity, SST variability, sea ice variability, and projected habitat suitability for the Greenland study region. Orange lines in each plot show 1,000 year smoothed mean values. The inset map shows the circumpolar region in dark blue and the region surrounding Greenland in red.

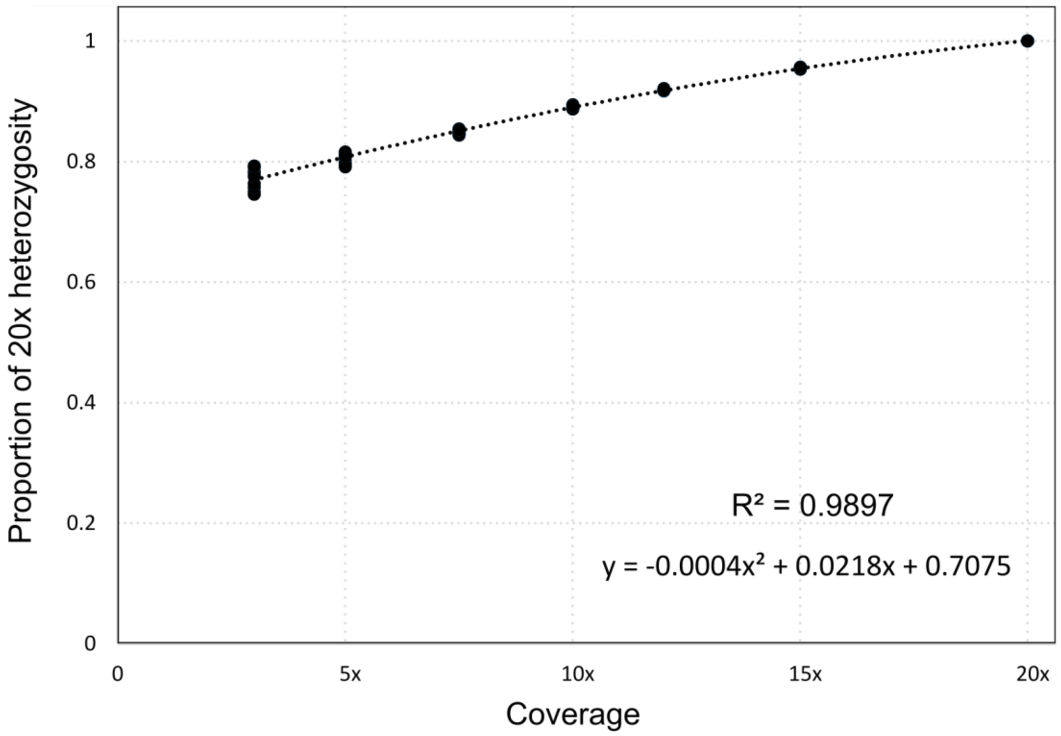


**Supplementary figure S10:** Polynomial regression line with an order of two calculated by downsampling a single high-coverage polar bear genome multiple times to correct for false negative heterozygous sites arising due to differences in coverage.

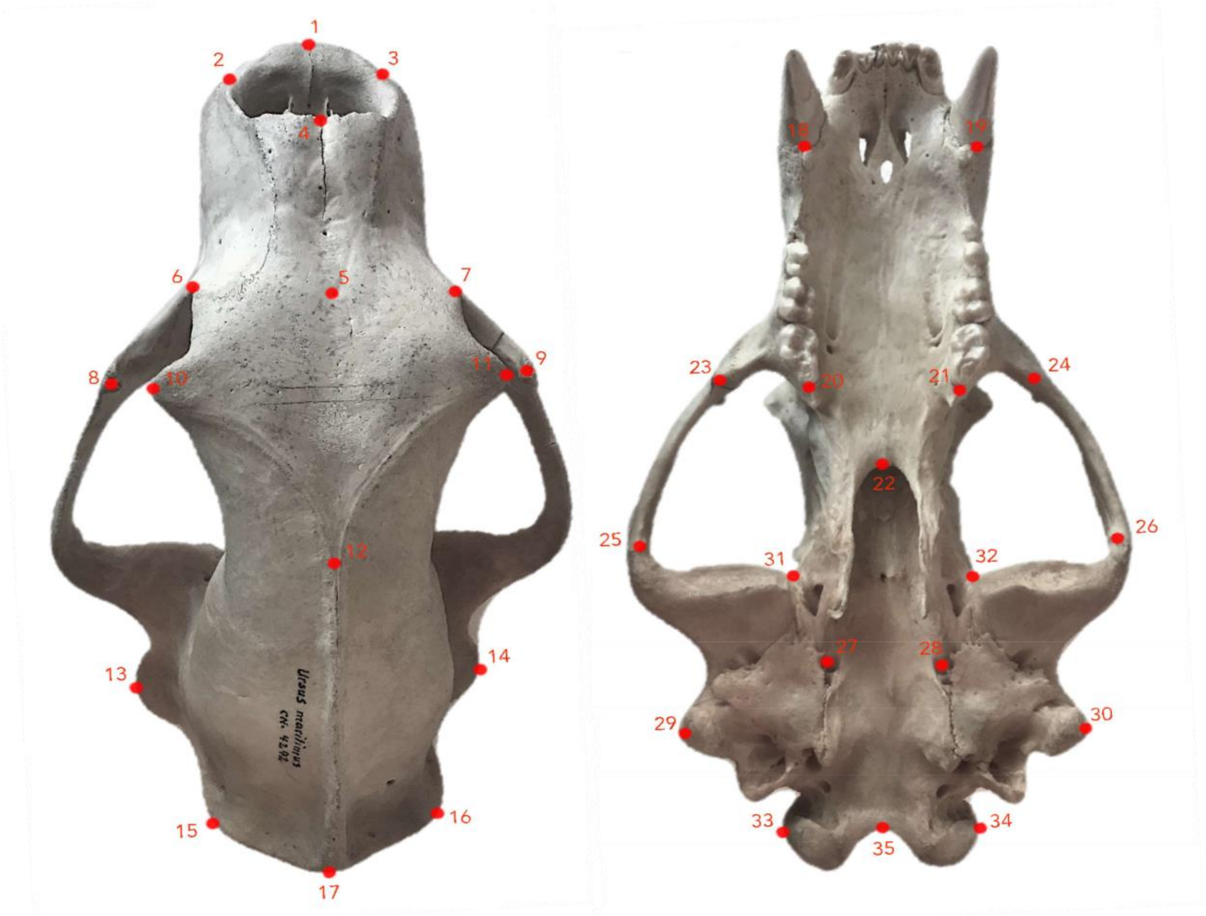


**Supplementary figure S11:** Dorsal and ventral view of the 35 landmarks considered in the geometric morphometric analyses. Landmarks 15, 16 and 17, were removed from all specimens, as many of the skulls were missing these landmarks. Each dot represents where the microscribe registered the coordinate.

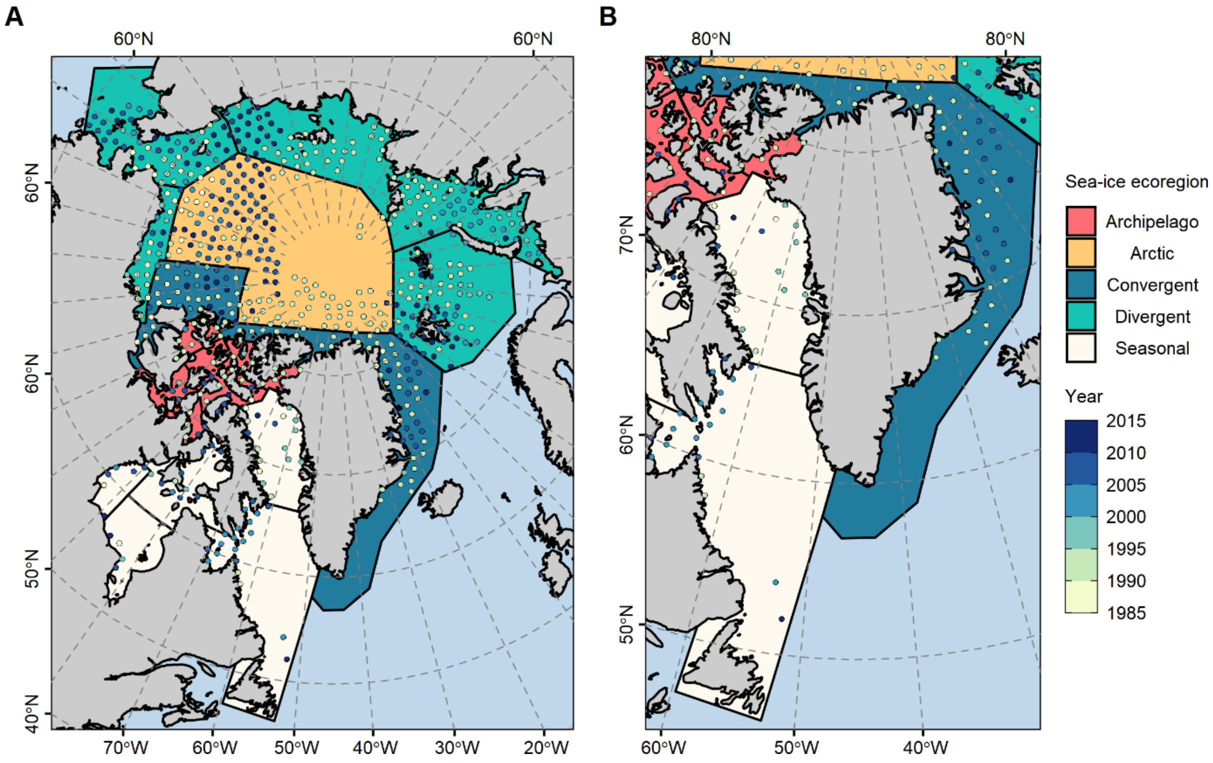


**Supplementary figure S12:** Map of occurrence records used for modelling polar bear habitat suitability. (A) Circumpolar distribution of the spatially thinned records used for building the habitat suitability model. (B) Records within the boundaries of the study region. Points are coloured by year of occurrence, with polygons coloured by polar bear ecoregion (18).

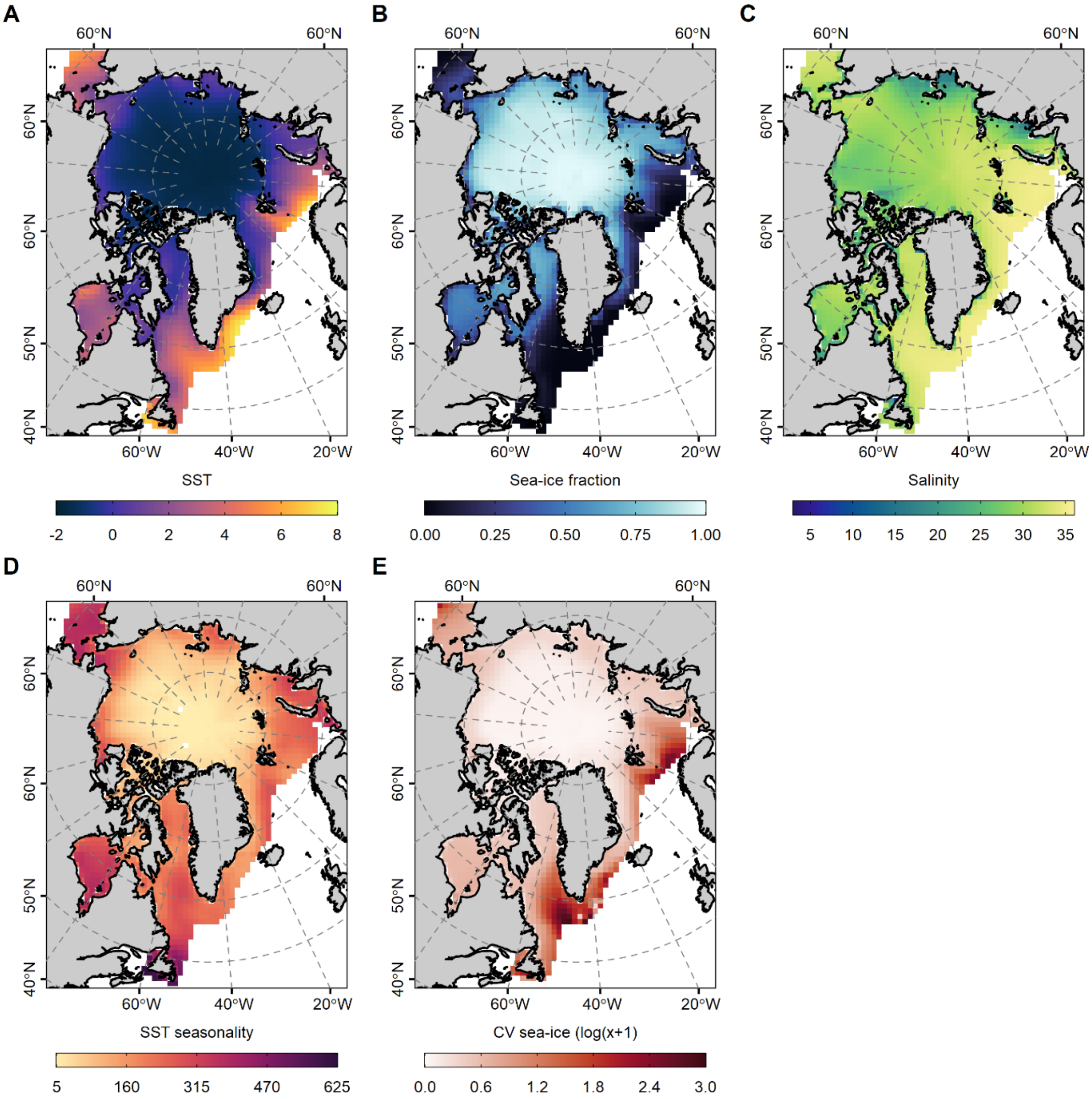


**Supplementary figure S13:** Climatological averages of temperature and environmental variables used in niche modelling. (A) Average sea-surface temperature, (B) average annual sea ice fraction, (C) average annual sea-surface salinity, (D) sea-surface temperature seasonality, (E) coefficient of variation in sea ice fraction. All variables were averaged over a 30-year period from 1985 – 2014 C.E. Note that E has been transformed for plotting purposes only.

**Supplementary text**

**Description of landmarks used for geometric morphometric analysis**

1. Inter Premaxillary suture at the alveolar margin
2. Maxillary-premaxillary suture at the left alveolar margin
3. Maxillary-premaxillary suture at the right alveolar margin
4. Most anterior edge of the nasal bones
5. Naso-frontal suture in the midline
6. Anteriormost point of the left orbital border
7. Anteriormost point of the right orbital border
8. Left, dorsal tip of the frontal process of the zygomatic arch
9. Right, dorsal tip of the frontal process of the zygomatic arch
10. Highest point of the left postorbital process
11. Highest point of the right postorbital process
12. Origin of sagittal crest
13. Highest point of left Squamosal
14. Highest point of right Squamosal
15. Highest point of left parietal bone
16. Highest point of right parietal bone
17. Most postero-ventral point of the occipital crest
18. Alveolar margin at the posterior aspect of the left canine
19. Alveolar margin at the posterior aspect of the right canine
20. The end point of left second molar
21. The end point of right second molar
22. Most posterior point of palatine bones suture
23. Lowest point on the anterior end of the left zygomatic arch
24. Lowest point on the anterior end of the right zygomatic arch
25. The left midpoint between squamosal and jugal suture
26. The right midpoint between squamosal and jugal suture
27. Left lateral border of suture between the basisphenoid and basioccipital
28. Right lateral border of suture between the basisphenoid and basioccipital
29. Highest point of the left squamosal
30. Highest point of the right squamosal
31. Anteriormost point of the left mandibular fossa
32. Anteriormost point of the right mandibular fossa
33. Left side of the occipital condyle
34. Right side of the occipital condyle
35. Most rostral point of the occipital condyle

**Supplementary video S1: Reconstruction of spatiotemporal habitat suitability for polar bears over the last 11,000 years.** Animation shows the relative stability in habitat suitability for the polar bear as predicted by our MaxEnt model, for the period 11,000 years ago to 2,000 C.E for the region surrounding Greenland. Can be downloaded from https://drive.google.com/file/d/1QoQ33N3KFfFlxdnS_HYkN-Itqz0A5y42/view?usp=sharing
