## Supplementary tables for "Impact of Holocene environmental change on the evolutionary ecology of an Arctic top predator"

| Sample ID | Biosample accession | Geographic region | Country/Region #2 | Polar bear management unit | Total number of hits (excluding | Estimated coverage from |
|---|---|---|---|---|---|---|
| Umar_MS392 | SAMN02045558 | Canada | Cornwallis Island | Lancaster sound | 95.365.141 | 4,70 |
| Umar_WH_JC004 | SAMN02045553 | Canada | West Hudson Bay | Western Hudson Bay | 85.840.346 | 4,39 |
| Umar_WH_JC002 | SAMN02045552 | Canada | West Hudson Bay | Western Hudson Bay | 95.218.724 | 4,81 |
| BGI-polarbear-PB_6 | SAMN02261835 | East Greenland | Ittoqqortoormiit | East Greenland | 57.747.863 | 1,80 |
| BGI-polarbear-PB_87 | SAMN02261874 | East Greenland | | East Greenland | 72.457.299 | 2,72 |
| BGI-polarbear-PB_2 | SAMN02261850 | East Greenland | Ittoqqortoormiit | East Greenland | 74.474.136 | 2,80 |
| BGI-polarbear-PB_8 | SAMN02261849 | East Greenland | Ittoqqortoormiit | East Greenland | 77.263.321 | 2,90 |
| BGI-polarbear-PB_9 | SAMN02261848 | East Greenland | Ittoqqortoormiit | East Greenland | 80.025.404 | 3,01 |
| BGI-polarbear-PB_31 | SAMN02261864 | East Greenland | Ittoqqortoormiit | East Greenland | 160.529.729 | 3,29 |
| BGI-polarbear-PB_83 | SAMN02261879 | East Greenland | | East Greenland | 90.034.546 | 3,38 |
| BGI-polarbear-PB_01 | SAMN02261873 | East Greenland | | East Greenland | 100.136.761 | 3,76 |
| BGI-polarbear-PB_86 | SAMN02261875 | East Greenland | | East Greenland | 106.513.085 | 3,99 |
| D24105 | SAMN16454158 | East Greenland | Northeast Greenland | East Greenland | 59.725.871 | 4,09 |
| BGI-polarbear-PB_29 | SAMN02261820 | East Greenland | Ittoqqortoormiit | East Greenland | 109.643.815 | 4,12 |
| BGI-polarbear-PB_19 | SAMN02261813 | East Greenland | Ittoqqortoormiit | East Greenland | 110.266.784 | 4,13 |
| BGI-polarbear-PB_90 | SAMN02261876 | East Greenland | | East Greenland | 110.979.774 | 4,16 |
| BGI-polarbear-PB_5 | SAMN02261843 | East Greenland | Ittoqqortoormiit | East Greenland | 111.219.573 | 4,17 |
| BGI-polarbear-PB_26 | SAMN02261817 | East Greenland | Ittoqqortoormiit | East Greenland | 112.621.349 | 4,22 |
| BGI-polarbear-PB_32 | SAMN02261823 | East Greenland | Ittoqqortoormiit | East Greenland | 113.818.041 | 4,28 |
| D24084 | SAMN16454146 | East Greenland | Northeast Greenland | East Greenland | 59.510.500 | 4,31 |
| BGI-polarbear-PB_27 | SAMN02261818 | East Greenland | Ittoqqortoormiit | East Greenland | 116.106.957 | 4,35 |
| BGI-polarbear-PB_37 | SAMN02261806 | East Greenland | Ittoqqortoormiit | East Greenland | 115.692.272 | 4,35 |
| BGI-polarbear-PB_4 | SAMN02261809 | East Greenland | Ittoqqortoormiit | East Greenland | 118.517.861 | 4,46 |
| BGI-polarbear-PB_38 | SAMN02261807 | East Greenland | Ittoqqortoormiit | East Greenland | 119.939.867 | 4,51 |
| BGI-polarbear-PB_35 | SAMN02261825 | East Greenland | Ittoqqortoormiit | East Greenland | 120.192.898 | 4,52 |
| BGI-polarbear-PB_30 | SAMN02261822 | East Greenland | Ittoqqortoormiit | East Greenland | 121.153.982 | 4,55 |
| BGI-polarbear-PB_34 | SAMN02261824 | East Greenland | Ittoqqortoormiit | East Greenland | 121.576.071 | 4,57 |
| BGI-polarbear-PB_39 | SAMN02261808 | East Greenland | Ittoqqortoormiit | East Greenland | 124.503.792 | 4,68 |
| BGI-polarbear-PB_40 | SAMN02261810 | East Greenland | Ittoqqortoormiit | East Greenland | 146.431.539 | 5,49 |
| D24109 | SAMN16454160 | East Greenland | Northeast Greenland | East Greenland | 80.902.456 | 5,56 |
| BGI-polarbear-PB_25 | SAMN02261816 | East Greenland | Ittoqqortoormiit | East Greenland | 202.449.886 | 7,59 |
| BGI-polarbear-PB_36 | SAMN02261805 | East Greenland | Ittorqortoormiut | East Greenland | 494.224.901 | 20,15 |
| BGI-polarbear-PB_33 | SAMN02261858 | East Greenland | Ittorqortoormiut | East Greenland | 563.458.134 | 21,03 |
| D24082 | SAMN16454145 | East Greenland | Northeast Greenland | East Greenland | 420.695.457 | 26,58 |
| BGI-polarbear-PB_89 | SAMN02261878 | East Greenland | | East Greenland | 690.668.750 | 28,49 |
| BGI-polarbear-PB_7 | SAMN02261826 | East Greenland | Ittorqortoormiut | East Greenland | 717.108.780 | 29,34 |
| BGI-polarbear-PB_3 | SAMN02261821 | East Greenland | Ittorqortoormiut | East Greenland | 727.256.341 | 29,88 |
| D24033 | SAMN10734296 | East Greenland | Northeast Greenland | East Greenland | 481.502.880 | 30,48 |
| BGI-polarbear-PB_28 | SAMN02261819 | East Greenland | Ittorqortoormiut | East Greenland | 750.615.339 | 30,86 |
| D24080 | SAMN16454143 | East Greenland | Northeast Greenland | East Greenland | 519.369.185 | 32,82 |
| PB12_N26030 | SAMN01057669 | Svalbard | Spitsbergen | Barents sea | 103.100.415 | 4,19 |
| PB17_N26024 | SAMN01057674 | Svalbard | Spitsbergen | Barents sea | 125.970.808 | 5,13 |
| PB16_N23797 | SAMN01057673 | Svalbard | Spitsbergen | Barents sea | 126.813.295 | 5,16 |
| PB18_N26025 | SAMN01057675 | Svalbard | Spitsbergen | Barents sea | 129.913.750 | 5,28 |
| PB15_N23694 | SAMN01057672 | Svalbard | Spitsbergen | Barents sea | 135.416.171 | 5,48 |
| PB13_N23355 | SAMN01057670 | Svalbard | Spitsbergen | Barents sea | 138.236.703 | 5,58 |
| PB5_N23949 | SAMN01057662 | Svalbard | Spitsbergen | Barents sea | 169.756.559 | 6,98 |
| PB11_N26029 | SAMN01057668 | Svalbard | Spitsbergen | Barents sea | 179.934.314 | 7,38 |
| PB14_N23379 | SAMN01057671 | Svalbard | Spitsbergen | Barents sea | 210.075.164 | 8,46 |
| PB1_N23531 | SAMN01057636 | Svalbard | Spitsbergen | Barents sea | 317.566.026 | 13,08 |
| PB9_N23985 | SAMN01057666 | Svalbard | Spitsbergen | Barents sea | 317.952.753 | 13,11 |
| PB10_N23997 | SAMN01057667 | Svalbard | Spitsbergen | Barents sea | 324.765.683 | 13,40 |
| PB6_N26028 | SAMN01057663 | Svalbard | Spitsbergen | Barents sea | 326.306.143 | 13,40 |
| PB2_N23604 | SAMN01057659 | Svalbard | Spitsbergen | Barents sea | 325.230.684 | 13,52 |
| PB4_N23917 | SAMN01057661 | Svalbard | Spitsbergen | Barents sea | 334.884.301 | 13,75 |
| PB8_N7968 | SAMN01057665 | Svalbard | Spitsbergen | Barents sea | 346.569.974 | 14,22 |
| PB3_N23719 | SAMN01057660 | Svalbard | Spitsbergen | Barents sea | 391.234.544 | 16,08 |
| PB7_N7773 | SAMN01057664 | Svalbard | Spitsbergen | Barents sea | 2.405.155.238 | 114,12 |
| BGI-polarbear-PB_56 | SAMN02261832 | West Greenland | Qaanaaq | Baffin Bay | 55.668.915 | 1,74 |
| BGI-polarbear-PB_61 | SAMN02261836 | West Greenland | Qaanaaq | Baffin Bay | 55.975.406 | 1,75 |
| BGI-polarbear-PB_71 | SAMN02261827 | West Greenland | Qaanaaq | Baffin Bay | 57.506.755 | 1,80 |
| BGI-polarbear-PB_50 | SAMN02261802 | West Greenland | Qaanaaq | Baffin Bay | 62.069.052 | 1,94 |
| BGI-polarbear-PB_60 | SAMN02261860 | West Greenland | Qaanaaq | Baffin Bay | 125.951.635 | 2,58 |
| BGI-polarbear-PB_24 | SAMN02261861 | West Greenland | Qaanaaq | Baffin Bay | 138.175.535 | 2,84 |
| BGI-polarbear-PB_49 | SAMN02261863 | West Greenland | Qaanaaq | Baffin Bay | 141.983.008 | 2,91 |
| BGI-polarbear-PB_46 | SAMN02261846 | West Greenland | Qaanaaq | Baffin Bay | 79.430.108 | 2,98 |
| BGI-polarbear-PB_51 | SAMN02261855 | West Greenland | Qaanaaq | Baffin Bay | 145.841.763 | 2,99 |
| BGI-polarbear-PB_106 | SAMN02261872 | West Greenland | Disko West | Baffin Bay | 82.783.862 | 3,11 |
| BGI-polarbear-PB_48 | SAMN02261842 | West Greenland | Qaanaaq | Baffin Bay | 101.911.353 | 3,82 |
| BGI-polarbear-PB_58 | SAMN02261834 | West Greenland | Qaanaaq | Baffin Bay | 136.571.838 | 4,80 |
| BGI-polarbear-PB_42 | SAMN02261870 | West Greenland | Qaanaq | Baffin Bay | 634.821.965 | 24,10 |
| BGI-polarbear-PB_22 | SAMN02261871 | West Greenland | Qaanaq | Baffin Bay | 666.567.654 | 24,94 |
| BGI-polarbear-PB_45 | SAMN02261868 | West Greenland | Qaanaq | Baffin Bay | 671.158.436 | 25,66 |
| BGI-polarbear-PB_47 | SAMN02261854 | West Greenland | Qaanaq | Baffin Bay | 695.216.602 | 26,44 |
| BGI-polarbear-PB_43 | SAMN02261811 | West Greenland | Qaanaq | Baffin Bay | 709.633.838 | 29,15 |
| BGI-polarbear-PB_105 | SAMN02261880 | West Greenland | Disko West | Baffin Bay | 747.577.226 | 30,90 |
| BGI-polarbear-PB_59 | SAMN02261839 | West Greenland | Avanersuaq | Kane Basin | 54.876.129 | 1,72 |
| BGI-polarbear-PB_66 | SAMN02261838 | West Greenland | Qaanaaq | Kane Basin | 56.513.403 | 1,77 |
| BGI-polarbear-PB_55 | SAMN02261831 | West Greenland | Avanersuaq | Kane Basin | 56.412.665 | 1,77 |
| BGI-polarbear-PB_74 | SAMN02261828 | West Greenland | Qaanaaq | Kane Basin | 57.161.320 | 1,79 |
| BGI-polarbear-PB_64 | SAMN02261837 | West Greenland | Qaanaaq | Kane Basin | 57.374.361 | 1,79 |
| BGI-polarbear-PB_57 | SAMN02261833 | West Greenland | Qaanaaq | Kane Basin | 58.446.043 | 1,83 |
| BGI-polarbear-PB_78 | SAMN02261830 | West Greenland | Qaanaaq | Kane Basin | 58.765.499 | 1,84 |
| BGI-polarbear-PB_53 | SAMN02261804 | West Greenland | Avanersuaq | Kane Basin | 59.805.211 | 1,87 |
| BGI-polarbear-PB_75 | SAMN02261829 | West Greenland | Qaanaaq | Kane Basin | 61.256.370 | 1,92 |
| BGI-polarbear-PB_52 | SAMN02261803 | West Greenland | Avanersuaq | Kane Basin | 61.584.358 | 1,93 |
| BGI-polarbear-PB_17 | SAMN02261866 | West Greenland | Avanersuaq | Kane Basin | 123.049.596 | 2,53 |
| BGI-polarbear-PB_81 | SAMN02261847 | West Greenland | Qaanaaq | Kane Basin | 68.436.188 | 2,57 |
| BGI-polarbear-PB_69 | SAMN02261862 | West Greenland | Qaanaaq | Kane Basin | 129.559.443 | 2,66 |
| BGI-polarbear-PB_13 | SAMN02261857 | West Greenland | Qaanaaq | Kane Basin | 131.256.752 | 2,69 |
| BGI-polarbear-PB_76 | SAMN02261869 | West Greenland | Qaanaaq | Kane Basin | 140.107.561 | 2,88 |
| BGI-polarbear-PB_11 | SAMN02261859 | West Greenland | Qaanaaq | Kane Basin | 142.025.989 | 2,91 |
| BGI-polarbear-PB_15 | SAMN02261852 | West Greenland | Avanersuaq | Kane Basin | 150.863.665 | 3,09 |
| BGI-polarbear-PB_18 | SAMN02261812 | West Greenland | Avanersuaq | Kane Basin | 86.007.265 | 3,22 |
| BGI-polarbear-PB_20 | SAMN02261814 | West Greenland | Avanersuaq | Kane Basin | 103.797.858 | 3,89 |
| BGI-polarbear-PB_54 | SAMN02261844 | West Greenland | Avanersuaq | Kane Basin | 108.356.205 | 4,07 |
| BGI-polarbear-PB_44 | SAMN02261841 | West Greenland | Qaanaaq | Kane Basin | 110.830.951 | 4,16 |
| BGI-polarbear-PB_72 | SAMN02261840 | West Greenland | Qaanaq | Kane Basin | 638.473.326 | 26,29 |
| BGI-polarbear-PB_79 | SAMN02261853 | West Greenland | Qaanaq | Kane Basin | 716.474.139 | 27,44 |
| BGI-polarbear-PB_62 | SAMN02261851 | West Greenland | Qaanaq | Kane Basin | 729.578.553 | 27,70 |
| BGI-polarbear-PB_10 | SAMN02261856 | West Greenland | Qaanaq | Kane Basin | 739.581.620 | 28,36 |
| BGI-polarbear-PB_68 | SAMN02261845 | West Greenland | Qaanaq | Kane Basin | 744.373.233 | 30,74 |
| BGI-polarbear-PB_12 | SAMN02261865 | West Greenland | Qaanaq | Kane Basin | 816.859.569 | 31,01 |
| BGI-polarbear-PB_14 | SAMN02261867 | West Greenland | | Unknown | 147.358.705 | 3,02 |
| BGI-polarbear-PB_104 | SAMN02261877 | West Greenland | | Unknown | 99.767.930 | 3,74 |
| BGI-polarbear-PB_21 | SAMN02261815 | West Greenland | Qaanaaq | Unknown | 113.619.631 | 4,25 |
| Giant_Panda | | | | | 1.768.461.133 | 31,40 |
| Spectacled bear | | | | | 899.678.196 | 39,91 |

| Sample ID | Geographic region | Management unit | Mean adjusted heterozygosity | Lower 5% | Upper 5% |
|---|---|---|---|---|---|
| BGI-polarbear-PB_10 | West Greenland | Kane Basin | 0,00083 | 0,00077 | 0,00090 |
| BGI-polarbear-PB_105 | West Greenland | Baffin Bay | 0,00080 | 0,00073 | 0,00086 |
| BGI-polarbear-PB_12 | West Greenland | Kane Basin | 0,00084 | 0,00078 | 0,00091 |
| BGI-polarbear-PB_22 | West Greenland | Baffin Bay | 0,00082 | 0,00076 | 0,00089 |
| BGI-polarbear-PB_42 | West Greenland | Baffin Bay | 0,00082 | 0,00075 | 0,00088 |
| BGI-polarbear-PB_43 | West Greenland | Baffin Bay | 0,00082 | 0,00075 | 0,00088 |
| BGI-polarbear-PB_45 | West Greenland | Baffin Bay | 0,00083 | 0,00076 | 0,00089 |
| BGI-polarbear-PB_47 | West Greenland | Baffin Bay | 0,00087 | 0,00080 | 0,00093 |
| BGI-polarbear-PB_62 | West Greenland | Kane Basin | 0,00083 | 0,00076 | 0,00089 |
| BGI-polarbear-PB_68 | West Greenland | Kane Basin | 0,00084 | 0,00077 | 0,00090 |
| BGI-polarbear-PB_72 | West Greenland | Kane Basin | 0,00084 | 0,00077 | 0,00090 |
| BGI-polarbear-PB_79 | West Greenland | Kane Basin | 0,00083 | 0,00077 | 0,00090 |
| BGI-polarbear-PB_25 | East Greenland | East Greenland | 0,00084 | 0,00076 | 0,00092 |
| BGI-polarbear-PB_28 | East Greenland | East Greenland | 0,00074 | 0,00068 | 0,00080 |
| BGI-polarbear-PB_3 | East Greenland | East Greenland | 0,00081 | 0,00075 | 0,00088 |
| BGI-polarbear-PB_33 | East Greenland | East Greenland | 0,00083 | 0,00077 | 0,00090 |
| BGI-polarbear-PB_36 | East Greenland | East Greenland | 0,00079 | 0,00073 | 0,00085 |
| BGI-polarbear-PB_40 | East Greenland | East Greenland | 0,00080 | 0,00071 | 0,00088 |
| BGI-polarbear-PB_7 | East Greenland | East Greenland | 0,00081 | 0,00074 | 0,00087 |
| BGI-polarbear-PB_89 | East Greenland | East Greenland | 0,00083 | 0,00077 | 0,00090 |
| D24082 | East Greenland | East Greenland | 0,00060 | 0,00055 | 0,00066 |
| D24080 | East Greenland | East Greenland | 0,00082 | 0,00075 | 0,00089 |
| D24109 | East Greenland | East Greenland | 0,00072 | 0,00064 | 0,00081 |
| D24033 | East Greenland | East Greenland | 0,00068 | 0,00062 | 0,00074 |

| LAB ID (Trent University) | Cat no (NHMD) /Museum ID | Geographic region | Locality/site | Polar bear management unit | Taxon | Year of sampling | Sex | $\delta^{13}C_{VPDB}$ | $\delta$ 13CSuess | $\delta^{15}N_{AIR}$ | wt%C | wt%N | C:N |
|---|---|---|---|---|---|---|---|---|---|---|---|---|---|
| 10366 | 692 | East Greenland | Dove Bugt | | Bearded seal | 1939 | F | -15,96 | -15,81 | +17,21 | 36,22 | 13,17 | 3,21 |
| 10367 | 828 | East Greenland | Zachenbergbugten | | Bearded seal | 1949 | F | -15,87 | -15,67 | +16,74 | 31,25 | 11,56 | 3,15 |
| 10372 | 797 | West Greenland | Godhavn | | Bearded seal | 1949 | F | -14,23 | -14,03 | +16,77 | 39,59 | 13,95 | 3,31 |
| 10363 | 622 | East Greenland | Scoresbysund | | Bearded seal | 1927 | M | -15,76 | -15,65 | +17,01 | 39,37 | 14,18 | 3,24 |
| 10365 | 687 | East Greenland | Kangerlussuaq | | Bearded seal | 1938 | M | -14,76 | -14,61 | +14,58 | 39,91 | 14,34 | 3,24 |
| 10373 | 831 | West Greenland | Godhavn | | Bearded seal | 1949 | M | -13,53 | -13,32 | +16,69 | 37,17 | 13,51 | 3,21 |
| 10374 | 1158 | West Greenland | Jakobshavn (Ilulissat) | | Bearded seal | 1919 | M | -14,47 | -14,38 | +16,59 | 35,78 | 12,80 | 3,26 |
| 10362 | 319 | East Greenland | Danmarks Havn | | Bearded seal | 1906 | | -14,94 | -14,87 | +15,74 | 32,90 | 12,05 | 3,18 |
| 10364 | 623 | East Greenland | Scoresbysund | | Bearded seal | 1928 | | -15,81 | -15,70 | +17,80 | 39,67 | 14,39 | 3,22 |
| 10368 | 1197 | East Greenland | | | Bearded seal | 1935 | | -13,60 | -13,46 | +14,95 | 39,37 | 14,23 | 3,23 |
| 10370 | 30 | West Greenland | Kangamiut | | Bearded seal | 1850 | | -13,36 | -13,34 | +16,32 | 40,84 | 14,98 | 3,18 |
| 10371 | 34 | West Greenland | Egedesminde | | Bearded seal | 1862 | | -13,35 | -13,33 | +17,21 | 38,10 | 13,80 | 3,22 |
| 10329 | 615 | East Greenland | Scoresbysund | | Harp seal | 1928 | F | -16,46 | -16,34 | +13,76 | 43,76 | 15,82 | 3,23 |
| 10332 | 620 | East Greenland | Scoresbysund | | Harp seal | 1928 | F | -15,54 | -15,42 | +14,97 | 38,13 | 13,77 | 3,23 |
| 10340 | 480 | West Greenland | | | Harp seal | 1948 | F | -14,41 | -14,22 | +16,36 | 39,62 | 14,37 | 3,22 |
| 10344 | 962 | West Greenland | Saqaq, Umanak | | Harp seal | 1963 | F | -15,20 | -14,90 | +16,34 | 38,28 | 13,64 | 3,27 |
| 10326 | 612 | East Greenland | Scoresbysund | | Harp seal | 1928 | M | -16,14 | -16,03 | +15,16 | 40,77 | 14,80 | 3,21 |
| 10328 | 614 | East Greenland | Scoresbysund | | Harp seal | 1928 | M | -16,74 | -16,62 | +14,06 | 39,54 | 14,37 | 3,21 |
| 10330 | 616 | East Greenland | Scoresbysund | | Harp seal | 1928 | M | -15,12 | -15,00 | +16,42 | 30,97 | 11,18 | 3,23 |
| 10331 | 617 | East Greenland | Scoresbysund | | Harp seal | 1927 | M | -15,65 | -15,54 | +15,02 | 36,05 | 13,12 | 3,20 |
| 10333 | 621 | East Greenland | Scoresbysund | | Harp seal | 1928 | M | -15,20 | -15,08 | +15,97 | 41,74 | 15,22 | 3,20 |
| 10334 | 852 | East Greenland | Scoresbysund | | Harp seal | 1928 | M | -16,96 | -16,84 | +13,66 | 38,62 | 14,09 | 3,20 |
| 10335 | 47 | West Greenland | Fiskenæs | | Harp seal | pre-1841 | M | -14,70 | -14,70 | +15,67 | 38,66 | 14,04 | 3,21 |
| 10336 | 56 | West Greenland | Fiskenæs | | Harp seal | pre-1811 | M | -14,24 | -14,24 | +15,97 | 35,12 | 12,86 | 3,18 |
| 10337 | 371 | West Greenland | Egedesminde | | Harp seal | pre-1911 | M | -14,57 | -14,57 | +17,49 | 38,22 | 13,76 | 3,24 |
| 10338 | 372 | West Greenland | Egedesminde | | Harp seal | pre-1911 | M | -14,27 | -14,27 | +16,31 | 44,23 | 16,01 | 3,22 |
| 10339 | 782 | West Greenland | Sarqaq, Nugsuak | | Harp seal | 1948 | M | -14,63 | -14,44 | +17,14 | 43,85 | 15,76 | 3,24 |
| 10342 | 960 | West Greenland | Saqaq, Umanak | | Harp seal | 1963 | M | -15,86 | -15,57 | +17,05 | 38,24 | 13,30 | 3,35 |
| 10343 | 961 | West Greenland | Saqaq, Umanak | | Harp seal | 1961 | M | -15,31 | -15,03 | +17,39 | 38,44 | 13,73 | 3,26 |
| 10327 | 613 | East Greenland | Scoresbysund | | Harp seal | 1927 | | -16,98 | -16,87 | +13,41 | 40,49 | 14,62 | 3,23 |
| 10325 | CN 1195 | East Greenland | Angmagsalik | | Harp seal | 1936 | | -15,42 | -15,28 | +15,65 | 39,99 | 14,45 | 3,23 |
| 10341 | 848 | West Greenland | Sarqaq | | Harp seal | 1949 | | -15,51 | -15,31 | +16,33 | 38,63 | 13,87 | 3,25 |
| 10379 | M 1267 | East Greenland | Kap København, Pearyland | | Hooded seal | 1983 | F | -15,01 | -14,50 | +18,98 | 37,28 | 13,30 | 3,27 |
| 10383 | 399 | West Greenland | Kronprinsens Eiland, Disko bugt | | Hooded seal | 1913 | F | -14,68 | -14,61 | +15,44 | 38,55 | 13,52 | 3,33 |
| 10378 | CN 1203 | East Greenland | Angmagsalik | | Hooded seal | 1935 | M | -15,33 | -15,19 | +16,45 | 34,98 | 12,34 | 3,31 |
| 10381 | 378 | West Greenland | Frederikshåb | | Hooded seal | pre-1912 | M | -12,76 | -12,76 | +18,22 | 40,14 | 14,45 | 3,24 |
| 10375 | 402 | East Greenland | | | Hooded seal | pre-1917 | | -14,67 | -14,67 | +15,94 | 33,76 | 12,02 | 3,27 |
| 10376 | CN 624 | East Greenland | Scoresbysund | | Hooded seal | 1927-1928 | | -15,64 | -15,64 | +18,04 | 40,86 | 14,54 | 3,28 |
| 10380 | 323 | West Greenland | Sukkertoppen | | Hooded seal | pre- 1909 | | -14,12 | -14,12 | +16,30 | 32,06 | 11,32 | 3,30 |
| 5254 | M 10746 | East Greenland | Ittorqortoqormiut | East Greenland | Polar bear | 2012 | F | -17,19 | -16,08 | +19,33 | 34,33 | 12,87 | 3,11 |
| 5255 | M 10747 | East Greenland | Ittorqortoqormiut | East Greenland | Polar bear | 2012 | F | -16,87 | -15,75 | +18,78 | 33,52 | 12,60 | 3,10 |
| 5256 | M 10754 | East Greenland | Ittoqqortoormiit | East Greenland | Polar bear | 2013 | F | -16,04 | -14,90 | +18,60 | 31,81 | 12,02 | 3,09 |
| 5257 | M 10814 | East Greenland | N-E Greenland | East Greenland | Polar bear | 2015 | F | -16,37 | -15,16 | +19,06 | 33,64 | 12,59 | 3,12 |
| 5258 | M 10819 | East Greenland | N-E Greenland | East Greenland | Polar bear | 2015 | F | -15,75 | -14,54 | +18,16 | 32,65 | 12,29 | 3,10 |
| 5251 | M 8612 | East Greenland | Kap Tobin, | East Greenland | Polar bear | 2001 | F | -16,24 | -15,41 | +17,97 | 32,35 | 11,74 | 3,21 |
| 5252 | M 8626 | East Greenland | Turner Ø, | East Greenland | Polar bear | 2001 | F | -16,51 | -15,68 | +18,96 | 32,44 | 11,96 | 3,16 |
| 5253 | M 8627 | East Greenland | Rømer Fjord, | East Greenland | Polar bear | 2001 | F | -15,87 | -15,04 | +18,78 | 32,90 | 12,39 | 3,10 |
| 5121 | M 10728 | West Greenland | N-W Greenland | Baffin Bay | Polar bear | 2011 | F | -15,21 | -14,13 | +20,59 | 33,14 | 12,32 | 3,14 |
| 5122 | M 10732 | West Greenland | N-W Greenland | Baffin Bay | Polar bear | 2011 | F | -15,52 | -14,44 | +20,99 | 37,23 | 13,53 | 3,21 |
| 5123 | M 10734 | West Greenland | N-W Greenland | Baffin Bay | Polar bear | 2011 | F | -15,27 | -14,19 | +21,18 | 34,58 | 12,84 | 3,14 |
| 5119 | M 8668 | West Greenland | Melville Bugten | Baffin Bay | Polar bear | 2006 | F | -14,12 | -13,18 | +21,54 | 33,38 | 12,43 | 3,13 |
| 5120 | M 10154 | West Greenland | Avanersuaq, NW | Kane Basin | Polar bear | 2009 | F | -15,54 | -14,52 | +23,26 | 38,09 | 14,00 | 3,17 |
| 5124 | M 10763 | West Greenland | Avernersuaq, NW | Kane Basin | Polar bear | 2013 | F | -15,57 | -14,43 | +21,47 | 33,00 | 12,29 | 3,13 |
| 5223 | M 10813 | East Greenland | N-E Greenland | East Greenland | Polar bear | 2015 | M | -16,31 | -15,10 | +19,08 | 33,08 | 12,44 | 3,10 |
| 5224 | M 10817 | East Greenland | N-E Greenland | East Greenland | Polar bear | 2015 | M | -16,04 | -14,83 | +19,87 | 34,09 | 12,75 | 3,12 |
| 5225 | M 10820 | East Greenland | N-E Greenland | East Greenland | Polar bear | 2015 | M | -16,22 | -15,01 | +20,34 | 33,77 | 12,60 | 3,13 |
| 5226 | M 10821 | East Greenland | N-E Greenland | East Greenland | Polar bear | 2015 | M | -16,16 | -14,95 | +18,94 | 33,31 | 12,54 | 3,10 |
| 5227 | M 10904 | East Greenland | N-E Greenland | East Greenland | Polar bear | 2014 | M | -16,06 | -14,88 | +19,65 | 32,14 | 12,08 | 3,10 |
| 5170 | M 8549 | East Greenland | Turner Ø, | East Greenland | Polar bear | 2001 | M | -15,75 | -14,93 | +18,87 | 33,88 | 12,69 | 3,11 |
| 5171 | M 8562 | East Greenland | Kap Brewster, | East Greenland | Polar bear | 2001 | M | -15,72 | -14,90 | +19,54 | 36,78 | 13,68 | 3,14 |
| 5172 | M 8599 | East Greenland | Manby Halvø, | East Greenland | Polar bear | 2001 | M | -15,63 | -14,81 | +18,96 | 33,69 | 12,64 | 3,11 |
| 5173 | M 8605 | East Greenland | Rømer Fjord, | East Greenland | Polar bear | 2001 | M | -16,10 | -15,27 | +20,62 | 33,25 | 12,39 | 3,13 |
| 5174 | M 8609 | East Greenland | Turner Ø, | East Greenland | Polar bear | 2001 | M | -15,51 | -14,69 | +19,01 | 34,47 | 12,92 | 3,11 |
| 5137 | M 10138 | West Greenland | N-W Greenland | Baffin Bay | Polar bear | 2009 | M | -16,34 | -15,31 | +19,62 | 31,08 | 11,58 | 3,13 |
| 5139 | M 10726 | West Greenland | N-W Greenland | Baffin Bay | Polar bear | 2011 | M | -15,10 | -14,02 | +22,09 | 34,84 | 12,90 | 3,15 |
| 5138 | M 10151 | West Greenland | Avanersuaq, NW | Kane Basin | Polar bear | 2009 | M | -15,30 | -14,27 | +21,62 | 33,39 | 12,42 | 3,13 |
| 5133 | M 8864 | West Greenland | Qaanaaq | Kane Basin | Polar bear | 2006 | M | -15,26 | -14,32 | +20,47 | 34,80 | 12,87 | 3,15 |
| 5134 | M 9758 | West Greenland | Thule. | Kane Basin | Polar bear | 2012 | M | -15,32 | -14,21 | +21,75 | 35,66 | 13,08 | 3,18 |
| 5135 | M 9759 | West Greenland | Thule. | Kane Basin | Polar bear | 2012 | M | -15,44 | -14,33 | +21,94 | 34,01 | 12,60 | 3,15 |
| 5136 | M 9760 | West Greenland | Thule. | Kane Basin | Polar bear | 2012 | M | -15,20 | -14,09 | +21,78 | 33,75 | 12,66 | 3,11 |
| 10270 | 560 | East Greenland | | | Ringed seal | 1928 | F | -16,87 | -16,76 | +15,04 | 37,15 | 13,22 | 3,28 |
| 10272 | 574 | East Greenland | | | Ringed seal | 1927 | F | -16,59 | -16,48 | +17,47 | 40,22 | 14,65 | 3,20 |
| 10274 | 591 | East Greenland | | | Ringed seal | 1928 | F | -16,86 | -16,75 | +15,33 | 39,07 | 14,20 | 3,21 |
| 10275 | 592 | East Greenland | | | Ringed seal | 1928 | F | -16,41 | -16,29 | +14,89 | 38,08 | 14,00 | 3,17 |
| 10278 | 597 | East Greenland | | | Ringed seal | 1928 | F | -16,61 | -16,49 | +17,66 | 40,50 | 14,76 | 3,20 |
| 10267 | M1237 | East Greenland | | | Ringed seal | 1994 | F | -16,71 | -16,03 | +14,82 | 40,10 | 14,72 | 3,18 |
| 10283 | 386 | West Greenland | | | Ringed seal | pre-1915 | F | -15,66 | -15,66 | +16,77 | 39,39 | 14,22 | 3,23 |
| 10284 | 486 | West Greenland | | | Ringed seal | 1924 | F | -14,27 | -14,17 | +17,84 | 38,91 | 14,30 | 3,17 |
| 10271 | 570 | East Greenland | | | Ringed seal | 1927 | M | -16,70 | -16,59 | +17,65 | 41,11 | 14,79 | 3,24 |
| 10273 | 581 | East Greenland | | | Ringed seal | 1928 | M | -16,55 | -16,43 | +17,68 | 38,26 | 13,97 | 3,20 |
| 10276 | 595 | East Greenland | | | Ringed seal | 1928 | M | -16,91 | -16,80 | +16,55 | 37,97 | 13,78 | 3,21 |
| 10277 | 596 | East Greenland | | | Ringed seal | 1928 | M | -16,62 | -16,50 | +12,60 | 41,59 | 15,17 | 3,20 |
| 10279 | 603 | East Greenland | | | Ringed seal | 1928 | M | -16,47 | -16,35 | +13,62 | 41,66 | 15,17 | 3,20 |
| 10280 | 606 | East Greenland | | | Ringed seal | 1928 | M | -16,55 | -16,43 | +14,62 | 37,05 | 13,49 | 3,20 |
| 10281 | 608 | East Greenland | | | Ringed seal | 1928 | M | -16,85 | -16,74 | +14,88 | 38,05 | 13,66 | 3,25 |
| 10268 | M1242 | East Greenland | | | Ringed seal | 1994 | M | -17,04 | -16,36 | +14,48 | 39,82 | 14,82 | 3,13 |
| 10269 | M1244 | East Greenland | | | Ringed seal | 1994 | M | -17,40 | -16,72 | +13,67 | 36,61 | 13,59 | 3,14 |
| 10285 | 781 | West Greenland | | | Ringed seal | 1948 | M | -14,38 | -14,19 | +18,87 | 42,46 | 15,45 | 3,21 |
| 10286 | 810 | West Greenland | | | Ringed seal | 1940 | M | -14,92 | -14,76 | +17,17 | 41,96 | 15,24 | 3,21 |
| 10282 | M1252 | West Greenland | | | Ringed seal | 1994 | M | -15,72 | -15,04 | +16,15 | 36,45 | 13,49 | 3,15 |
| 10287 | 713017 | West Greenland | Qaasuitsup | | Ringed seal | 1971 | | -14,23 | -13,87 | +16,79 | 39,96 | 14,19 | 3,28 |
| 10288 | 713031 | West Greenland | Qaasuitsup | | Ringed seal | 1971 | | -15,54 | -15,17 | +16,35 | 39,47 | 14,07 | 3,27 |
| 10289 | 713037 | West Greenland | Qaasuitsup | | Ringed seal | 1971 | | -15,30 | -14,93 | +17,89 | 38,94 | 14,05 | 3,23 |
| 10291 | 713080 | West Greenland | Qaasuitsup | | Ringed seal | 1971 | | -14,99 | -14,62 | +18,72 | 41,50 | 14,66 | 3,30 |
| 10292 | 713081 | West Greenland | Qaasuitsup | | Ringed seal | 1971 | | -15,32 | -14,95 | +18,75 | 34,97 | 12,51 | 3,26 |
| 10293 | 713082 | West Greenland | Qaasuitsup | | Ringed seal | 1971 | | -16,16 | -15,79 | +16,14 | 39,09 | 13,71 | 3,33 |
| 10294 | 713102 | West Greenland | Qaasuitsup | | Ringed seal | 1971 | | -15,48 | -15,11 | +16,63 | 40,16 | 14,29 | 3,28 |
| 10295 | 713110 | West Greenland | Qaasuitsup | | Ringed seal | 1971 | | -15,38 | -15,01 | +16,94 | 39,92 | 14,33 | 3,25 |
| 10296 | 713170 | West Greenland | Qaasuitsup | | Ringed seal | 1971 | | -16,20 | -15,83 | +17,10 | 36,21 | 12,93 | 3,27 |
| 10290 | 713176 | West Greenland | Qaasuitsup | | Ringed seal | 1971 | | -15,78 | -15,42 | +17,72 | 36,82 | 13,14 | 3,27 |
| 10300 | M11 CN 461 | East Greenland | | | Walrus | 1924 | F | -15,90 | -15,80 | +13,24 | 39,79 | 14,42 | 3,22 |
| 10301 | M11 CN 726 | East Greenland | | | Walrus | 1928 | F | -15,93 | -15,82 | +10,61 | 41,21 | 14,96 | 3,21 |
| 10303 | M11 CN 729 | East Greenland | | | Walrus | pre-1935 | F | -16,10 | -16,10 | +11,37 | 35,97 | 13,05 | 3,21 |
| 10304 | M11 CN 732 | East Greenland | | | Walrus | 1927 | F | -16,22 | -16,11 | +10,64 | 39,98 | 14,57 | 3,20 |
| 10312 | M11 CN 2 | West Greenland | | | Walrus | pre-1843 | F | -14,34 | -14,34 | +10,87 | 37,91 | 13,84 | 3,19 |
| 10313 | M11 CN 276 | West Greenland | | | Walrus | 1898 | F | -13,56 | -13,51 | +11,49 | 30,80 | 11,17 | 3,21 |
| 10320 | M11 CN 818 | West Greenland | | | Walrus | 1939 | F | -15,04 | -14,88 | +10,40 | 41,32 | 14,85 | 3,25 |
| 10321 | M11 CN 819 | West Greenland | | | Walrus | 1939 | F | -14,61 | -14,46 | +10,03 | 39,60 | 14,47 | 3,19 |
| 10324 | M11 CN 822 | West Greenland | | | Walrus | 1940 | F | -14,54 | -14,38 | +10,71 | 41,08 | 14,87 | 3,22 |
| 10305 | 34588 | East Greenland | Illorqortormiit | | Walrus | 1988 | M | -16,95 | -16,37 | +11,92 | 37,38 | 13,63 | 3,20 |
| 10306 | 34597 | East Greenland | Illorqortormiit | | Walrus | 1991 | M | -17,31 | -16,68 | +14,06 | 31,42 | 11,38 | 3,22 |
| 10307 | 34600 | East Greenland | Illorqortormiit | | Walrus | 1993 | M | -16,49 | -15,82 | +14,71 | 30,28 | 11,13 | 3,17 |
| 10308 | 34701 | East Greenland | Illorqortormiit | | Walrus | 1993 | M | -16,65 | -15,98 | +13,51 | 27,05 | 9,90 | 3,19 |
| 10309 | 34702 | East Greenland | Illorqortormiit | | Walrus | 1993 | M | -16,61 | -15,94 | +15,26 | 27,96 | 10,24 | 3,19 |
| 10310 | 34733 | East Greenland | Illorqortormiit | | Walrus | 1990 | M | -17,14 | -16,53 | +11,35 | 34,42 | 12,53 | 3,20 |
| 10311 | 34735 | East Greenland | Illorqortormiit | | Walrus | 1990 | M | -16,75 | -16,13 | +13,26 | 40,74 | 14,85 | 3,20 |
| 10302 | M11 CN 728 | East Greenland | | | Walrus | 1927 | M | -16,33 | -16,22 | +11,43 | 42,00 | 15,17 | 3,23 |
| 10315 | M11 CN 465 | West Greenland | | | Walrus | pre-1926 | M | -13,89 | -13,89 | +12,63 | 37,16 | 13,43 | 3,23 |
| 10318 | M11 CN 816 | West Greenland | | | Walrus | 1939 | M | -14,92 | -14,76 | +10,61 | 39,43 | 14,35 | 3,20 |
| 10319 | M11 CN 817 | West Greenland | | | Walrus | 1939 | M | -14,96 | -14,80 | +10,32 | 40,33 | 14,64 | 3,21 |
| 10322 | M11 CN 820 | West Greenland | | | Walrus | 1940 | M | -14,60 | -14,44 | +10,84 | 35,55 | 12,90 | 3,21 |
| 10323 | M11 CN 821 | West Greenland | | | Walrus | 1940 | M | -14,99 | -14,83 | +10,04 | 32,03 | 11,66 | 3,20 |
| 10297 | M11 CN 290 | East Greenland | | | Walrus | pre-1909 | | -16,04 | -16,04 | +12,81 | 39,10 | 14,24 | 3,20 |
| 10298 | M11 CN 291 | East Greenland | | | Walrus | pre-1909 | | -15,88 | -15,88 | +11,85 | 37,37 | 13,64 | 3,19 |
| 10299 | M11 CN 293 | East Greenland | | | Walrus | pre-1909 | | -16,15 | -16,15 | +13,24 | 37,84 | 13,74 | 3,21 |
| 10314 | M11 CN 403 | West Greenland | | | Walrus | pre-1915 | | -14,56 | -14,56 | +10,45 | 40,19 | 14,50 | 3,23 |
| 10316 | M11 CN 477 | West Greenland | | | Walrus | pre-1924 | | -13,85 | -13,85 | +11,77 | 37,44 | 13,76 | 3,17 |
| 10317 | M11 CN 478 | West Greenland | | | Walrus | pre-1924 | | -14,50 | -14,50 | +13,33 | 42,28 | 15,35 | 3,21 |
| 14152 | NHMO-DMA-29601/1-O | West Greenland | Sukkertoppen | | Bearded Seal | 1892 | | -15,21 | -15,21 | 17,33 | 41,48 | 15,02 | 3,22 |
| 14166 | NHMO-DMA-32440/1-O | West Greenland | Sukkertoppen | | Bearded Seal | 1894 | | -13,29 | -13,29 | 17,5 | 43,2 | 14,8 | 3,4 |
| 14161 | NHMO-DMA-32999/1-O | West Greenland | Godthåb | | Bearded Seal | 1892 | | -14,05 | -14,05 | 14,62 | 42,19 | 15,35 | 3,2 |
| 14158 | NHMO-DMA-32435/1-O | West Greenland | Sukkertoppen | | Hooded Seal | 1894 | | -14,22 | -14,22 | 16,19 | 39,9 | 13,63 | 3,41 |
| 14167 | NHMO-DMA-32436/1-O | West Greenland | Sukkertoppen | | Hooded Seal | 1894 | | -14,99 | -14,99 | 14,74 | 42,6 | 14,4 | 3,45 |
| 14165 | NHMO-DMA-32437/1-O | West Greenland | Sukkertoppen | | Hooded Seal | 1894 | | -16,07 | -16,07 | 15,13 | 43,7 | 14,5 | 3,52 |
| 14146 | NHMO-DMA-32968/1-O | West Greenland | Sukkertoppen | | Hooded Seal | 1893 | | -14,11 | -14,11 | 16,11 | 39,42 | 13,85 | 3,32 |
| 14144 | NHMO-DMA-32973/1-O | East Greenland | Danmarkstredet | | Hooded Seal | 1897 | | -15,28 | -15,28 | 12,55 | 42,19 | 14,15 | 3,48 |
| 14157 | NHMO-DMA-32990/1-O | East Greenland | Danmarkstredet | | Hooded Seal | 1897 | M | -14,54 | -14,54 | 15,49 | 40,73 | 14,16 | 3,35 |
| 14154 | NHMO-DMA-32994/1-O | East Greenland | Danmarkstredet | | Hooded Seal | 1897 | F | -15,54 | -15,54 | 14,29 | 40,66 | 14,59 | 3,25 |
| 14147 | NHMO-DMA-33013/1-O | East Greenland | Danmarkstredet | | Hooded Seal | 1888 | | -15,7 | -15,7 | 16,2 | 41 | 14,11 | 3,39 |
| 14162 | NHMO-DMA-32969/1-O | West Greenland | Sukkertoppen | | Hooded Seal | 1893 | | -14,73 | -14,73 | 14,77 | 41,6 | 14,8 | 3,28 |
| 14149 | NHMO-DMA-32992/1-O | East Greenland | Danmarkstredet | | Hooded Seal | 1888 | | -15,23 | -15,23 | 13,79 | 40,5 | 14,27 | 3,31 |

| Cat no (NHMD) | Field no (AU) | Geographic region | Locality | Polar bear | Year of sampling | Sex | Age (years) |
|---|---|---|---|---|---|---|---|
| CN5394 | | East Greenland | Angmagssalik | East Greenland | 1966 | M | 13 |
| M8650 | 26393 | East Greenland | Barclay Bay | East Greenland | 2003 | F | 8 |
| CN5438 | | East Greenland | Danmarks Havn, N-E Greenland | East Greenland | 1969-70 | M | 13 |
| M8570 | 21727 | East Greenland | Danneborg | East Greenland | 2000 | F | 8 |
| M8585 | 21749 | East Greenland | Danneborg | East Greenland | 2000 | F | 5 |
| M9755 | 33933 | East Greenland | East Greenland | East Greenland | 2007 | M | 13 |
| M9752 | 33939 | East Greenland | East Greenland | East Greenland | 2007 | F | 9 |
| M9745 | 33955 | East Greenland | East Greenland | East Greenland | 2007 | F | 19 |
| M9734 | 33957 | East Greenland | East Greenland | East Greenland | 2007 | M | 16 |
| M8561 | 21711 | East Greenland | Henry Land | East Greenland | 2000 | M | 8 |
| M8639 | 26376 | East Greenland | Hurry Fjord | East Greenland | 2004 | M | 6 |
| CN5355 | | East Greenland | Ikagteq, Sermilik Fjord | East Greenland | 1967 | F | 6 |
| M8672 | 33927 | East Greenland | Ittoqqortoormiit | East Greenland | 2006 | M | 12 |
| M8670 | 33929 | East Greenland | Ittoqqortoormiit | East Greenland | 2006 | M | 9 |
| M8676 | 33931 | East Greenland | Ittoqqortoormiit | East Greenland | 2006 | M | 8 |
| M8675 | 33934 | East Greenland | Ittoqqortoormiit | East Greenland | 2006 | F | 5 |
| M9740 | 33935 | East Greenland | Ittoqqortoormiit | East Greenland | 2008 | F | 9 |
| M8666 | 33937 | East Greenland | Ittoqqortoormiit | East Greenland | 2006 | M | 17 |
| M8659 | 33938 | East Greenland | Ittoqqortoormiit | East Greenland | 2006 | F | 7 |
| M8667 | 33947 | East Greenland | Ittoqqortoormiit | East Greenland | 2006 | F | 15 |
| M8674 | 33951 | East Greenland | Ittoqqortoormiit | East Greenland | 2006 | M | 6 |
| M8673 | 33952 | East Greenland | Ittoqqortoormiit | East Greenland | 2006 | F | 8 |
| M9747 | 33954 | East Greenland | Ittoqqortoormiit | East Greenland | 2007 | M | 9 |
| M10778 | 39469 | East Greenland | Ittoqqortoormiit | East Greenland | 2010 | M | 7 |
| M10779 | 39470 | East Greenland | Ittoqqortoormiit | East Greenland | 2010 | F | 5 |
| M10780 | 39471 | East Greenland | Ittoqqortoormiit | East Greenland | 2010 | M | 8 |
| M10781 | 39472 | East Greenland | Ittoqqortoormiit | East Greenland | 2010 | M | 6 |
| M10783 | 39474 | East Greenland | Ittoqqortoormiit | East Greenland | 2010 | F | 6 |
| M10787 | 39488 | East Greenland | Ittoqqortoormiit | East Greenland | 2010 | F | 17 |
| M9751 | 39710 | East Greenland | Ittoqqortoormiit | East Greenland | 2008 | M | 6 |
| M9757 | 39717 | East Greenland | Ittoqqortoormiit | East Greenland | 2008 | F | 16 |
| M9756 | 39718 | East Greenland | Ittoqqortoormiit | East Greenland | 2008 | M | 9 |
| M9742 | 39729 | East Greenland | Ittoqqortoormiit | East Greenland | 2009 | M | 6 |
| M10139 | 39858 | East Greenland | Ittoqqortoormiit | East Greenland | 2009 | M | 12 |
| M10142 | 39861 | East Greenland | Ittoqqortoormiit | East Greenland | 2009 | M | 13 |
| M10144 | 39863 | East Greenland | Ittoqqortoormiit | East Greenland | 2009 | M | 10 |
| M10722 | 39865 | East Greenland | Ittoqqortoormiit | East Greenland | 2009 | M | 13 |
| M10148 | 39872 | East Greenland | Ittoqqortoormiit | East Greenland | 2009 | M | 6 |
| M10149 | 39873 | East Greenland | Ittoqqortoormiit | East Greenland | 2009 | M | 13 |
| M10744 | 46752 | East Greenland | Ittoqqortoormiit | East Greenland | 2012 | M | 10 |
| M10745 | 46753 | East Greenland | Ittoqqortoormiit | East Greenland | 2012 | M | 6 |
| M10746 | 46754 | East Greenland | Ittoqqortoormiit | East Greenland | 2012 | F | 7 |
| M10747 | 46755 | East Greenland | Ittoqqortoormiit | East Greenland | 2012 | F | 5 |
| M10749 | 46757 | East Greenland | Ittoqqortoormiit | East Greenland | 2012 | M | 7 |
| M10754 | 48703 | East Greenland | Ittoqqortoormiit | East Greenland | 2013 | F | 13 |
| M10756 | 48705 | East Greenland | Ittoqqortoormiit | East Greenland | 2013 | M | 8 |
| M10757 | 48706 | East Greenland | Ittoqqortoormiit | East Greenland | 2013 | M | 14 |
| M10758 | 48707 | East Greenland | Ittoqqortoormiit | East Greenland | 2013 | M | 9 |
| CN5359 | | East Greenland | Ittoqqortoormiit | East Greenland | 1967 | F | 10 |
| CN5358 | | East Greenland | Ittoqqortoormiit | East Greenland | 1967 | M | 14 |
| CN5360 | | East Greenland | Ittoqqortoormiit | East Greenland | 1967 | M | 17 |
| CN5367 | | East Greenland | Ittoqqortoormiit | East Greenland | 1967 | F | 25 |
| CN5368 | | East Greenland | Ittoqqortoormiit | East Greenland | 1967 | F | 14 |
| CN5366 | | East Greenland | Ittoqqortoormiit | East Greenland | 1967 | F | 16 |
| CN5347 | | East Greenland | Kangerdlussuaq | East Greenland | 1966 | F | 4 |
| CN5350 | | East Greenland | Kangerdlussuaq | East Greenland | 1966 | M | 9 |
| M8554 | 21695 | East Greenland | Kap Brewster | East Greenland | 2000 | F | 16 |
| M8562 | 21712 | East Greenland | Kap Brewster | East Greenland | 2001 | M | 7 |
| M8576 | 21737 | East Greenland | Kap Brewster | East Greenland | 1999 | M | 8 |
| M8619 | 22209 | East Greenland | Kap Brewster | East Greenland | 2000 | F | 19 |
| CN5353 | | East Greenland | Kap Hammer | East Greenland | 1966 | F | 8 |
| M8600 | 21772 | East Greenland | Kap Hope/Kap Tobin | East Greenland | 2001 | F | 20 |
| M8630 | 22227 | East Greenland | Kap Russel | East Greenland | 2000 | M | 28 |
| M8537 | 21668 | East Greenland | Kap Swainson | East Greenland | 2001 | M | 8 |
| M8545 | 21678 | East Greenland | Kap Swainson | East Greenland | 2000 | F | 5 |
| M8547 | 21680 | East Greenland | Kap Swainson | East Greenland | 2000 | F | 22 |
| M8641 | 26879 | East Greenland | Kap Sweinson | East Greenland | 2004 | F | 21 |
| M8553 | 21693 | East Greenland | Kap Tobin | East Greenland | 1999 | F | 10 |
| M8558 | 21705 | East Greenland | Kap Tobin | East Greenland | 1999 | M | 23 |
| M8566 | 21719 | East Greenland | Kap Tobin | East Greenland | 2000 | F | 5 |
| M8567 | 21720 | East Greenland | Kap Tobin | East Greenland | 2000 | F | 12 |
| M8577 | 21739 | East Greenland | Kap Tobin | East Greenland | 2000 | M | 16 |
| M8588 | 21755 | East Greenland | Kap Tobin | East Greenland | 2000 | F | 19 |
| M8598 | 21770 | East Greenland | Kap Tobin | East Greenland | 1999 | M | 9 |
| M8612 | 22201 | East Greenland | Kap Tobin | East Greenland | 2001 | F | 12 |
| M8651 | 26394 | East Greenland | Kap Tobin | East Greenland | 2004 | F | 27 |
| CN5380 | | East Greenland | Kiataq, SE Greenland | East Greenland | 1967 | F | 10 |
| CN5390 | | East Greenland | Kulusuk, SE Greenland | East Greenland | 1967 | M | 7 |
| M10896 | 35165 | East Greenland | N-E Greenland | East Greenland | 2014 | M | 13 |
| M10901 | 35166 | East Greenland | N-E Greenland | East Greenland | 2014 | M | 6 |
| M10897 | 35168 | East Greenland | N-E Greenland | East Greenland | 2014 | M | 11 |
| M10735 | 43101 | East Greenland | N-E Greenland | East Greenland | 2011 | M | 6 |
| M10740 | 43109 | East Greenland | N-E Greenland | East Greenland | 2011 | M | 10 |
| M10814 | 50413 | East Greenland | N-E Greenland | East Greenland | 2015 | F | |
| M10820 | 50416 | East Greenland | N-E Greenland | East Greenland | 2015 | M | |
| M10817 | 50419 | East Greenland | N-E Greenland | East Greenland | 2015 | M | |
| M10819 | 50420 | East Greenland | N-E Greenland | East Greenland | 2015 | F | |
| M10821 | 50421 | East Greenland | N-E Greenland | East Greenland | 2015 | M | |
| M10813 | 50422 | East Greenland | N-E Greenland | East Greenland | 2015 | M | |
| M8536 | 21666 | East Greenland | | East Greenland | 2000 | F | 14 |
| M8538 | 21669 | East Greenland | | East Greenland | 2000 | M | 10 |
| M8542 | 21675 | East Greenland | | East Greenland | 2000 | M | 9 |
| M8556 | 21703 | East Greenland | | East Greenland | 1999 | F | 5 |
| M8550 | | East Greenland | | East Greenland | 2000 | M | 10 |
| M8627 | 22221 | East Greenland | Rømer Fjord | East Greenland | 2001 | F | 6 |
| CN5397 | | East Greenland | Sermiligaq, Angmagssalik | East Greenland | 1966 | F | 10 |
| CN5396 | | East Greenland | Sermiligaq, Angmagssalik | East Greenland | 1966 | M | 22 |
| M8541 | 21674 | East Greenland | Stewart Ø | East Greenland | 2000 | F | 5 |
| CN5391 | | East Greenland | Tiniteqilaq, Angmagssalik | East Greenland | 1967 | F | 14 |
| M8572 | 21729 | East Greenland | Turner | East Greenland | 2000 | M | 7 |
| M8560 | 21710 | East Greenland | Turner Ø | East Greenland | 1999 | F | 6 |
| M8609 | 21792 | East Greenland | Turner Ø | East Greenland | 2001 | M | 8 |
| M8858 | 33921 | East Greenland | Unartoq, Ittoqqrtoormiit | East Greenland | 2006 | M | 6 |
| M8634 | 26370 | East Greenland | Unnartaai | East Greenland | 2004 | M | 9 |
| M8584 | 21748 | East Greenland | Vega Sund | East Greenland | 2000 | F | 5 |
| CN5352 | | East Greenland | Watkins fjord | East Greenland | 1966 | F | 12 |
| CN4453 | | West Greenland | "Devils thumb" Melville Bay | Baffin Bay | 1936 | M | 15-20 |
| M10151 | 42252 | West Greenland | Avanersuaq | Kane Basin | 2009 | M | |
| M10154 | 42256 | West Greenland | Avanersuaq | Kane Basin | 2009 | F | |
| M10763 | 48717 | West Greenland | Avernersuaq | Kane Basin | 2013 | F | 8 |
| CN5371 | | West Greenland | Herberts island, Thule | Kane Basin | 1967 | M | 5-7 |
| CN4109 | | West Greenland | Kane Basin | Kane Basin | 1940 | F | 15 |
| CN4105 | | West Greenland | Kane Basin | Kane Basin | 1940 | F | 8-9 |
| CN4106 | | West Greenland | Kane Basin | Kane Basin | 1940 | F | |
| CN4101 | | West Greenland | Kap. Sydv. (?) North Greenland | Kane Basin | 1940 | F | 10-12 |
| M8668 | | West Greenland | Melville Bugten | Baffin Bay | 2006 | F | |
| CN5452 | | West Greenland | N of Thoms Island, Melville Bay | Baffin Bay | 1972 | F | |
| M10138 | 39483 | West Greenland | N-W Greenland | Baffin Bay | 2009 | M | |
| M10726 | 41444 | West Greenland | N-W Greenland | Baffin Bay | 2011 | M | 15 |
| M10728 | 41447 | West Greenland | N-W Greenland | Baffin Bay | 2011 | F | 19 |
| M10732 | 41453 | West Greenland | N-W Greenland | Baffin Bay | 2011 | F | 24 |
| M10734 | 41455-2 | West Greenland | N-W Greenland | Baffin Bay | 2011 | F | 8 |
| M8864 | | West Greenland | Qaanaaq | Kane Basin | 2006 | M | |
| CN5374 | | West Greenland | Semerssuaq, Dundas | Kane Basin | 1968 | F | 9 |
| CN5413 | | West Greenland | Thule | Kane Basin | 1969 | M | |
| M9760 | 39720 | West Greenland | Thule. | Kane Basin | 2008 | M | 6 |
| M9759 | 39731 | West Greenland | Thule. | Kane Basin | 2009 | M | 10 |
| M9758 | 39732 | West Greenland | Thule. | Kane Basin | 2009 | M | |
| CN1355 | | West Greenland | Ubernivik | Baffin Bay | N/A | M | 7-8 |
| CN579 | | West Greenland | Upernavik | Baffin Bay | N/A | M | 10 |
| CN587 | | West Greenland | Upernivik | Baffin Bay | N/A | M | 6 |
| CN590 | | West Greenland | Upernivik | Baffin Bay | N/A | F | 6 |
| CN591 | | West Greenland | Upernivik | Baffin Bay | N/A | F | 20 |
| CN586 | | West Greenland | Upernivik | Baffin Bay | N/A | M | 8-9 |

| **Dataset** | **Source** |
| --- | --- |
| Incidental sightings of marine mammals | http://ipt.vliz.be/eurobis/resource?r=imr_mammals |
| Laidre et al (2018) | https://doi.org/10.1002/ece3.3809 |
| Laidre et al (2022) | https://doi.org/10.1126/science.abk2793 |
| Makinen & Vanhatalo (2018) | https://doi.org/10.1111/ddi.12776 |
| Peacock et al (2015) | https://doi.org/10.1371/journal.pone.0112021.s019 |
| GBIF - Polar bear observations | https://doi.org/10.15468/dl.5hhypa |
| GBIF - Arctic Species Trend Index | https://doi.org/10.15468/dl.9ev52f |
| Durner et al (2009) | https://doi.org/10.1890/07-2089.1 |
| Kovacs & Andersen (2010) | https://doi.org/10.21334/npolar.2010.246c1053 |
| Lydersen et al (2017) | https://doi.org/10.21334/npolar.2017.132248b4 |
| Hamilton et al (2019) | https://doi.org/10.21334/npolar.2019.e1cd54e1 |
| Rode et al (2016) | https://doi.org/10.5061/dryad.n5509 |
| Rode (2021) | https://doi.org/10.5066/P9KM5FT2 |
| Aerial Surveys of Arctic Marine Mammals | https://doi.org/10.7289/v51v5bzm |
| USGS - Polar bear tracking data (ARGOS locations) | https://www.usgs.gov/centers/asc/science/tracking-data-releases |
| USGS - Polar bear tracking data (1985-2015 Southern Beaufort Sea) | https://www.usgs.gov/centers/asc/science/tracking-data-releases |
